# Nup98-dependent transcriptional memory is established independently of transcription

**DOI:** 10.1101/2020.10.12.336073

**Authors:** Pau Pascual-Garcia, Shawn C. Little, Maya Capelson

## Abstract

Cellular ability to mount an enhanced transcriptional response upon repeated exposure to external cues has been termed transcriptional memory, which can be maintained epigenetically through cell divisions. The majority of mechanistic knowledge on transcriptional memory has been derived from bulk molecular assays, and this phenomenon has been found to depend on a nuclear pore component Nup98 in multiple species. To gain an alternative perspective on the mechanism and on the contribution of Nup98, we set out to examine single-cell population dynamics of transcriptional memory by monitoring transcriptional behavior of individual *Drosophila* cells upon initial and subsequent exposures to steroid hormone ecdysone. To this end, we combined single-molecule RNA FISH with mathematical modeling, and found that upon hormone exposure, cells rapidly activate a low-level transcriptional response, but simultaneously, begin a slow transitioning into a specialized memory state, characterized by a high rate of expression. Strikingly, our modeling predicted that this transition between non-memory and memory states is independent of the transcription stemming from initial activation, and we were able to confirm this prediction experimentally by showing that inhibiting transcription during initial ecdysone exposure did not interfere with memory establishment. Together, our findings reveal that Nup98’s role in transcriptional memory is to stabilize the forward rate of conversion from low to high expressing state, and that induced genes engage in two separate behaviors – transcription itself and the establishment of epigenetically propagated transcriptional memory.

## Introduction

Organisms need to adapt to varying changes continuously, and how cells adjust their transcriptional responses to different types of stimuli is critical for proper development and survival. One mechanism of cellular adaptation relies on the cells’ ability to build a more robust transcriptional response upon exposure to a signal that they have previously experienced. This phenomenon by which cells increase their re-activation kinetics, after a period of repression, is called transcriptional memory (Avramova, 2015; Bonifer and Cockerill, 2017; D’Urso and Brickner, 2017; Fabrizio et al., 2019). The ability of cells to increase their transcriptional response upon subsequent exposure (or “remember” their previous exposure) can be propagated through cell division, and is thus considered epigenetic. Transcriptional memory is well-characterized in genes that respond to environmental changes: for example, in budding yeast, rounds of inositol starvation trigger the transcriptional memory response for *INO1* (which encodes the enzyme catalyzing the limiting step in the biosynthesis of inositol) (Ahmed et al., 2010; Brickner et al., 2007; Light et al., 2010); or in plants, where pre-exposure to various abiotic stresses (e.g., drought, salinity) intensifies their resistance by elevating transcript levels from a subset of stress-response genes (Ding et al., 2013, 2012; Lämke et al., 2016; Liu et al., 2016; Sani et al., 2013). In both cases, transcriptional memory provides an advantage to survive environmental challenges by mounting a more robust transcriptional response that facilitates the adjustment to new conditions. Additionally, transcriptional memory is not only articulated through increased re-activation but also by transcriptional down-regulation of unnecessary genes. In yeast cells, repetitive carbon-source shifts produce not only a strong transcriptional re-activation of specific genes but also promote hyper-repression of unnecessary genes (Lee et al., 2018). Thus, organisms coordinate their transcriptional programs to prevent the loss of cellular energy and to optimize resources for a robust response that allows for increased fitness to changing environments.

Importantly, the primed state of transcription caused by the pre-exposure to a cue is not unique to environmental signals and can be found in different cellular processes. In mammalian cells, interferon-induced transcriptional memory is thought to confer an enhanced immune response to infectious agents (Gialitakis et al., 2010a; Kamada et al., 2018; Light et al., 2013), and in *Drosophila*, transcriptional memory has been reported in the context of signaling by steroid hormone ecdysone. Ecdysone or 20-hydroxyecdysone (20E) and its nuclear hormone receptor complex EcR/USP coordinate a transcriptional response that is essential for multiple events during fly development (Hill et al., 2013). A defined set of early ecdysone-induced genes, such as *ecdysone-induced protein 74EF (E74)* and *early gene at 23 (E23)*, are activated by the hormone receptor and remain primed for enhanced expression upon a second exposure to ecdysone, for at least 24 hours (Pascual-Garcia et al., 2017). Further evidence that transcriptional memory processes occur in organismal development comes from a study of re-activation dynamics during *Drosophila* embryogenesis, where transgenes were shown to re-activate more frequently and more rapidly after cell division if they were transcribing in the previous cell cycle (Ferraro et al., 2016).

Mechanistically, several models have been put forward to explain how transcriptional memory may work. One such model proposed that memory is based primarily on the cytoplasmic inheritance of a key regulatory factor, which is build up during the initial exposure and remains at high levels through the period of memory (Kundu and Peterson, 2009). Such a mode of inheritance regulates the long-term transcriptional memory of the yeast *GAL1* gene, which is repressed in normal glucose-containing media, but can be activated by switching cells to the alternative carbon source galactose (Brickner et al., 2007; Kundu et al., 2007; Zacharioudakis et al., 2007). In this case, the high expression of GAL1 regulators during initial activation and their subsequent continuous presence are thought to participate in the higher re-activation dynamics upon second exposure to galactose (Kundu and Peterson, 2010; Sood et al., 2017; Zacharioudakis et al., 2007). Another set of proposed models focus on the nuclear mode of inheritance of a transcriptional state, with chromatin and architectural features of the gene as the functional memory marks (D’Urso and Brickner, 2017; Kundu and Peterson, 2009; Randise-Hinchliff and Brickner, 2018). For instance, genes that display transcriptional memory have been shown to associate with a poised form of RNA Pol II pre-initiation complex (PIC) through a repression period, which is thought to allow faster re-activation kinetics (D’Urso et al., 2016; Light et al., 2013). This model of transcriptional memory centers on the binding of gene-specific transcriptional factors to cis-acting DNA elements and on contacts of the gene with nuclear pore complex (NPC) components (or Nups). These interactions can lead to an altered chromatin structure, characterized by changes in histone modifications and the acquisition of histone variants (Garvey Brickner et al., 2019; Kuhn and Capelson, 2019; Pascual-Garcia and Capelson, 2014; Sood and Brickner, 2014). Significant efforts have been made to understand how transcriptional memory operates for the yeast *INO1* gene, which is recruited to the NPC upon activation and remains anchored to the NPC after transcriptional shut off for several cell divisions. Recruitment of transcriptional factor Sfl1 at *INO1* promoter and contact with Nup100 (homolog of metazoan Nup98) have been shown to promote the incorporation of histone variant H2A.Z and H3K4me2, both of which have been linked to memory establishment in multiple systems (Bonifer et al., 2016; Brickner et al., 2007; D’Urso et al., 2016; Gialitakis et al., 2010b; Lämke et al., 2016; Light et al., 2013, 2010; Muramoto et al., 2010; Petter et al., 2011).

Studies from several model systems have revealed that Nup98 is an evolutionarily conserved factor necessary for transcriptional memory. In addition to INO1 signaling described above, interferon-induced genes require Nup98 for memory in human cells (Light et al., 2013). However, this memory event occurs at genes in the nuclear interior, which reflects the dynamic (shuttling on and off the pore) behavior of Nup98 in metazoan cells (Capelson et al., 2010; Kalverda et al., 2010). At both interferon-responsive genes and the *INO1* locus, Nup98 was found to be required for the deposition of the memory-associated H3K4Me2 mark (Light et al., 2013). Nup98 is similarly necessary for transcriptional memory in ecdysone-induced signaling. In *Drosophila* embryonic cells, loss of Nup98 does not affect transcription of ecdysone-induced genes during initial ecdysone exposure, but results in a poor memory response during re-induction conditions (Pascual-Garcia et al., 2017). At the same time, Nup98 was found to facilitate the maintenance of ecdysone-induced enhancer-promoter loops of genes like *E74* and *E23*, providing evidence that Nups can influence transcriptional memory through changes in chromatin contacts (Pascual-Garcia et al., 2017). This functional role of Nup98 in loop formation mimics similar findings in yeast, where NPC component Mlp1 is important for the intragenic loop formed between the promoter and 3’-end of galactose-inducible genes (Tan-Wong et al., 2009). Thus, influence of NPC components on transcriptional memory appears to be complex and reveals the intricate network of coordinated events that may have to occur to establish memory. Yet, although it is essential for transcriptional memory in multiple systems, exactly how Nup98 contributes to the enhanced transcriptional re-activation remains unclear. Investigation of this function of Nup98 provides an opportunity to understand the memory phenomenon further.

Previous work on transcriptional memory has primarily involved analysis of whole populations of cells, using bulk biochemical and molecular assays to investigate its mechanisms (Brickner et al., 2007; Ding et al., 2012; Gialitakis et al., 2010b; Pascual-Garcia et al., 2017). In this work, we aimed to understand how transcriptional memory could work at the level of single cells and how single-cell behaviors can give rise to population transcriptional outcomes. We took a combined approach based on precise quantification of ecdysone-induced *E74* mRNAs, single-molecule RNA FISH (smFISH), and mathematical modeling to describe transcriptional memory and the role of Nup98 from a cell population perspective. Comparing predicted population distributions, based on our modeling approaches, to actual distributions of nascent transcriptional states, obtained by smFISH, allowed us to define the transcriptional parameters that change in the reinduced vs. induced states. Similarly, we were able to determine which transcriptional parameters are regulated by Nup98, which helps understand the molecular contribution of Nup98 to the transcriptional process. We discovered that upon hormone induction, cells rapidly activate transcription, but slowly transition into a specialized memory state characterized by high *E74* expression. Strikingly, we found that transition into the memory state is triggered by ecdysone, but is nonetheless independent of the extent of transcriptional activity during the initial induction. Our results introduce a possible model that can account for cell population dynamics during transcriptional memory response, allow us to rule out some of the previously proposed models of transcriptional memory for the ecdysone-mediated memory, and suggest a functional role of Nup98 in driving the formation of a memory state that is separate from on-going transcription.

## Results

### 1. Assessing Nup98-dependent transcriptional memory in absolute units

We have previously shown that ecdysone-inducible genes exhibit transcriptional memory, and that Nup98 is required for the proper establishment and/or maintenance of the primed memory state (Pascual-Garcia et al., 2017). To further investigate the role of Nup98 in modulating transcription, we examined the transcription dynamics of the ecdysone-responsive gene *E74* during repeated hormone exposure in more detail. *Drosophila* S2 cells were exposed to synthetic 20E for a 4 hours initial induction, then washed and allowed to recover for 24 hours before re-exposure to 20E for an additional 4 hours (Figure 1A). We assayed *E74* expression dynamics during both exposures by collecting cells every 30 minutes, isolating mRNA, and performing RT-qPCR on control/dsWhite and Nup98 knockdown cells. To aid in quantitative analysis, we adapted a previously described protocol (Petkova et al., 2014) to estimate the absolute number of *E74* mRNA molecules per cell. We performed RT-qPCR using known numbers of *in vitro* transcribed RNAs as template spanning six orders of magnitude. This allowed us to construct a calibration curve relating the qPCR threshold cycle Ct to the number of input molecules (Figure Supplemental 1A; see Experimental Procedures). We measured the number of cells from which we extracted mRNA for each experimental condition and time point. We also estimated the mRNA recovery efficiency of the mRNA extraction step. Together, these measurements allowed us to convert experimental Ct values into absolute numbers of *E74* mRNA molecules per cell, generating a detailed comparison of *E74* expression dynamics during hormone exposure in control and Nup98 knockdown cells (Figure 1B).

**Figure 1.**
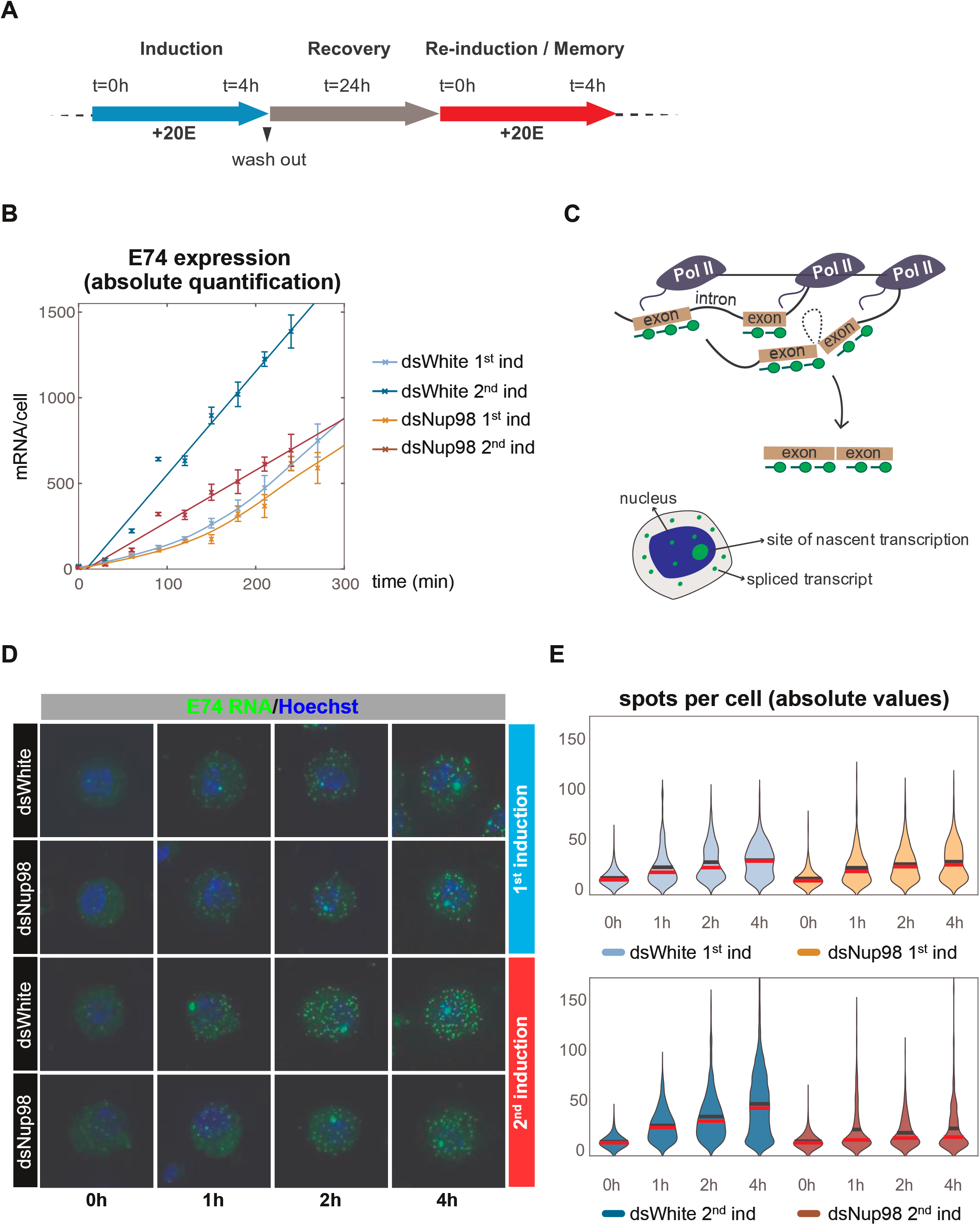
Absolute quantification of *E74* induction and transcriptional memory. **A**. Overview of ecdysone/20E treatment. Cells were initially treated with 5 μM of 20E, washed-out and recovered in fresh media for 24 hours. Memory response was assessed by incubating cells with 5 μM of 20E after recovery period. **B**. Absolute number of *E74* mRNAs per cell as a function of time after either the first or second induction in control (dsWhite) and Nup98 knockdown (dsNup98). Samples were collected every 30 min for both inductions. Error bars represent standard deviation of the mean of three experiments. **C**. Schematic of smFISH labeling of sites of nascent transcription and spliced transcripts in either the nucleus or cytoplasm. **D**. Representative images of *E74* expression during the first and second induction. **E**. Violin plots of *E74* puncta per cell. Mean and median indicated by black and red horizontal lines.

Using this approach, we found as expected, that the number of *E74* mRNA molecules per cell reaches considerably higher amounts during re-induction than during initial induction in control dsWhite-treated cells (Figure 1B). *E74* transcripts accumulate slowly during the first 2 hours of initial hormone exposure before increasing. Moreover, the accumulation trajectory in Nup98-depleted cells is not different from control during the first induction. In contrast, accumulation is rapidly onset during the second induction in control cells, whereas in Nup98 knockdown conditions, the second induction is more similar to the first induction (Figure 1B). This supports the proposed requirement for Nup98 in transcriptional memory, in agreement with previous work from our and other laboratories (Light et al., 2013; Pascual-Garcia et al., 2017). We verified that the efficiency of Nup98 depletion was similar during both the first and second inductions (Figure Supplemental 1B). This shows that the response of *E74* to initial exposure does not require normal levels of Nup98; instead, cells require Nup98 to rapidly re-express *E74* upon repeated exposure.

In the simplest model of hormone-induced transcription, all loci would be rapidly activated upon hormone treatment and then begin producing mRNAs at a constant rate. If true, then mRNA content should be nearly uniform between cells, particularly at later times after induction and at high expression rates observed during control second induction, when cells will have ample opportunity for their expression levels to closely approach the mean level. To ask whether *E74* expression is uniform between cells, and to gain insight into the nature of hormone-induced activity and memory response, we employed smFISH to measure *E74* expression in single cells. *E74* mRNAs were labeled with a set of 67 probes complimentary to exon sequences and imaged by confocal microscopy (Figure 1C). smFISH reveals two classes of labeled objects: relatively dim, diffraction-limited puncta enriched in the cytoplasm representing mature mRNAs, and one or more bright nuclear-localized puncta corresponding to sites of nascent transcription (Figures 1C, D). To estimate *E74* mRNA levels, we counted the number of mRNA puncta at 0, 1, 2, and 4 hours post-induction and observed that the average number of puncta per cell accumulated with dynamics that mirrored those obtained by absolute qPCR (Figure 1E). mRNA content of puncta also increased in a manner consistent with qPCR (Figure Supplemental 1C). As expected, during the first induction, dsWhite-treated and Nup98-depleted cells accumulated puncta at similar rates, while during the second induction, control cells accumulated mRNA puncta more rapidly and to higher levels than Nup98-depleted cells or cells undergoing the initial induction (Figure 1E). Importantly, we observed that *E74* expression is heterogeneous, with mRNA amounts exhibiting broad distribution at all times and under all induction conditions. Even under the most highly expressed condition at 4 hours after the second induction, 20% of cells possess 20 puncta or fewer, whereas <5% contain 100 or greater. These results suggest that *E74* activation occurs stochastically across the cell population, rather than occurring in a synchronized manner. Heterogeneous mRNA levels seen during the second induction further suggest that the role of Nup98 is to increase average expression levels without increasing the synchrony of the hormone response during the second induction.

### 2. Nup98 modulates *E74* mRNA production without affecting export or degradation

To further explore the role of Nup98 in the memory response, we undertook additional quantitative measurements of *E74* mRNA dynamics. Our preceding results (Figure 1) confirmed a Nup98-dependent increase in the rate of *E74* mRNA accumulation during the second induction. Generally, the rate of mRNA accumulation depends on both mRNA production and degradation. Degradation of mRNA may be affected by the process of mRNA export, which interfaces with NPCs (Rodríguez-Navarro and Hurt, 2011; Tutucci and Stutz, 2011). Certain NPC components, including, Nup98, have been previously implicated in mRNA export through interactions with the mRNA export machinery (Blevins et al., 2003; Chakraborty et al., 2006; Powers et al., 1997). mRNA export could in turn affect mRNA stability either directly or indirectly. Since prior work has not addressed whether Nup98 affects *E74* mRNA levels through these processes, we first asked whether Nup98 knockdown induces a change in *E74* mRNA trafficking out of the nucleus or in the rate of degradation of *E74*.

To study this scenario, we examined the smFISH images for evidence of altered nucleo-cytoplasmic transport, assessing the fraction of mRNA puncta that are found outside of the Hoechst-based mask, out of the total amount of mRNA puncta per cell (Supplemental Figure 2A). We found that this fraction does not change in control/dsWhite versus Nup98 knockdown cells in any of the four conditions we tested, with >85% of mRNAs found in the cytoplasm across conditions (Figure 2A). Next, we asked whether *E74* mRNA stability is altered upon Nup98 knockdown, for which we monitored *E74* mRNA levels following transcriptional arrest. We inhibited transcriptional elongation by treating cells with flavopiridol (FP), a potent inhibitor of p-TEFb, which phosphorylates Ser2 of the C-terminal domain of RNA Pol II (Chao and Price, 2001). *E74* mRNA synthesis was stimulated with 20E for 4 hours, after which the hormone was washed out and transcription blocked with FP. As a result of transcriptional blockage, the mRNA levels of *E74* mRNAs declined and we monitored them for 240 min, with time points collected every 30 min. We concluded that the degradation rates were largely unchanged between Nup98-depleted cells and control conditions, with *E74* mRNA lifetimes of 111 minutes and 117 minutes in Nup98-depleted and control cells, respectively (Figure 2B), which were determined by fitting to a first-order reaction. We concluded that neither mRNA export nor mRNA degradation are major contributors to the phenotype of Nup98 in transcriptional memory. Instead, our data suggests that the role of Nup98 in transcriptional memory is mediated through regulation of the transcriptional process, which is supported by the previously identified binding of Nup98 to the promoters and enhancers of ecdysone-inducible genes (Pascual-Garcia et al., 2017).

**Figure 2.**
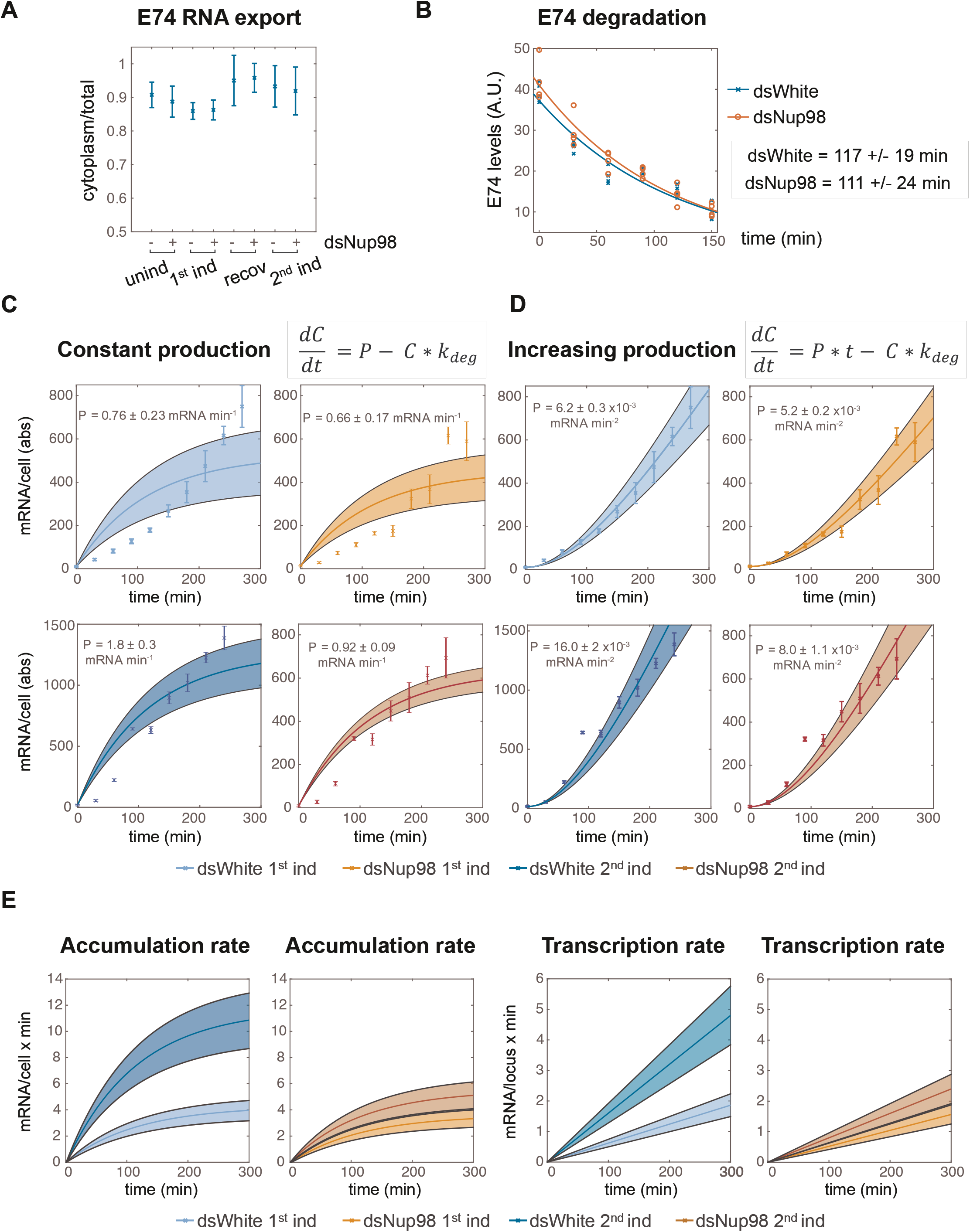
Altered Nup98 levels modulate transcription rates, not mRNA export or degradation. **A.** Fraction of *E74* mRNA found in cytoplasm, as assessed by smFISH, with Nup98 knockdown (+) or control knockdown (-) prior to the first induction (unind), after 4 hours (1^st^ ind), 24 hours after hormone removal (recov), and 4 hours after 2^nd^ induction. Error bars represent standard deviation of the mean. **B**. Degradation rates of *E74* mRNA assessed by qPCR. Cells were induced for 4 hours, washed-out and collected every 30 minutes after the addition of 1 μM flavopiridol at zero minutes. Experiment was repeated three times and mRNA lifetimes were determined by fit to exponential curves. **C-D**. *E74* transcription rates increase during 20E exposure. Data points with error bars obtained by qPCR as shown in Fig. 1B. Shaded bands indicate 95% confidence intervals for the fit rates P for the models indicated. Top row: first induction; second row: second induction. C: Best fit of data to model of constant transcription rates P. D: Best fit of data to model of time-dependent increasing *E74* transcription rate. **E**. Comparison of the rates of mRNA accumulation (left) and underlying transcription rates (right) between first and second induction in control and dsNup98 cells.

Our combined knowledge of the degradation rate along with the accumulation dynamics in absolute units allowed us to test quantitative models describing the effect on Nup98 transcription. In the simplest model, Nup98 would act to boost the average transcription rate from a low constant mRNA production rate during the first induction to a large constant rate during the second induction. To determine whether a model of constant production correctly describes the data, we found the best-fitting values for the production rates under each induction and knockdown combination (Figure 2C). In this straightforward model, the change in mRNA concentration per time equals mRNA production rate P minus the degradation rate, which was determined experimentally (Figure 2B). The estimate of the degradation rate permitted us to obtain best-fit values for P (Figure 2C). As expected, the value for production rate for the second induction in control cells was higher than during the first induction by a factor of 2.4x, whereas the production rates are similar in first induction control and Nup98 knockdown conditions under both exposures. However, constant production rates provide a poor description of the data (Figure 2C and Supplemental Figure 2B). In all cases, the constant production model predicts the fastest accumulation during the first two hours of expression, with gradual attenuation of the accumulation as transcript levels reach steady-state. The data instead exhibits a gradual and slow increase in the rate of accumulation during the majority of interval for hormone exposure. This is most obvious for the first induction in both control and Nup98 knockdown cells. The results indicate that the simplest explanation for Nup98 activity, boosting a constant production rate, is incompatible with observation.

Instead of constant production rate, the data suggested that the production of *E74* mRNA increases with time. We therefore fit a model in which production of new mRNAs increases linearly with time, and found that this model provided an excellent fit (Figure 2D and Supplemental Figure 2B). We observed that the production rates increase at similar rates in three scenarios: first induction in control, and both inductions in Nup98 knockdown conditions. In contrast, during the second induction in control cells, the production rate increases at more than twice the rate of the first induction. This can be clearly visualized by taking the derivative of the fit accumulation curves to reveal the accumulation rate, i.e., the change in the average number of mRNAs per cell as a function of time (Figure 2E). The accumulation rates are similar in control first induction and Nup98 depletion first and second induction, but become increasingly fast in control second induction. The accumulation rates begin to stabilize at around 4 mRNAs per minute during the first induction but reach greater than 10 mRNAs per minute during the second induction in control cells (Figure 2E, left). During hormone exposure, the transcription rate climbs continuously, reaching only about 2 mRNAs per minute per locus during the first induction but climbing to 5 mRNA per minute per locus during the second induction in control cells (Figure 2E, right). By the end of the fourth hour of the second exposure, this corresponds to an average production rate per locus similar to that observed for the most highly expressed genes in *Drosophila* embryos (Little et al., 2013; Zoller et al., 2018). Overall, these results show that Nup98 changes transcription of *E74* without affecting mRNA trafficking or stability. Moreover, normal levels of Nup98 are required not simply to promote a higher level of transcription. Instead, Nup98 appears to promote a continuous increase in transcription rates upon additional hormone exposure.

### 3. A two-state model for the mechanism of Nup98-dependent transcription rate increase

We sought to explore models describing the Nup98-dependent increase in the rate of mRNA production. We began with a two-state model in which loci switch between active and inactive states (Figure 3A-B). During the active state, new RNA Pol II molecules enter productive elongation at a rate given by k_Pol_. For simplicity, we assumed that while hormone is present, the rate switching into the active state k_A_ is much greater than the rate of switching into the inactive state k_-A_. This ensures that once active, loci are in essence irreversibly committed to the active state as long as the hormone is present. These assumptions are reasonable given the large but slow increase in transcript levels observed over the duration of the hormone exposure. In this model, the average production rate per locus is given by the loading rate k_Pol_ multiplied by the fraction of active loci (Figure 3B).

**Figure 3.**
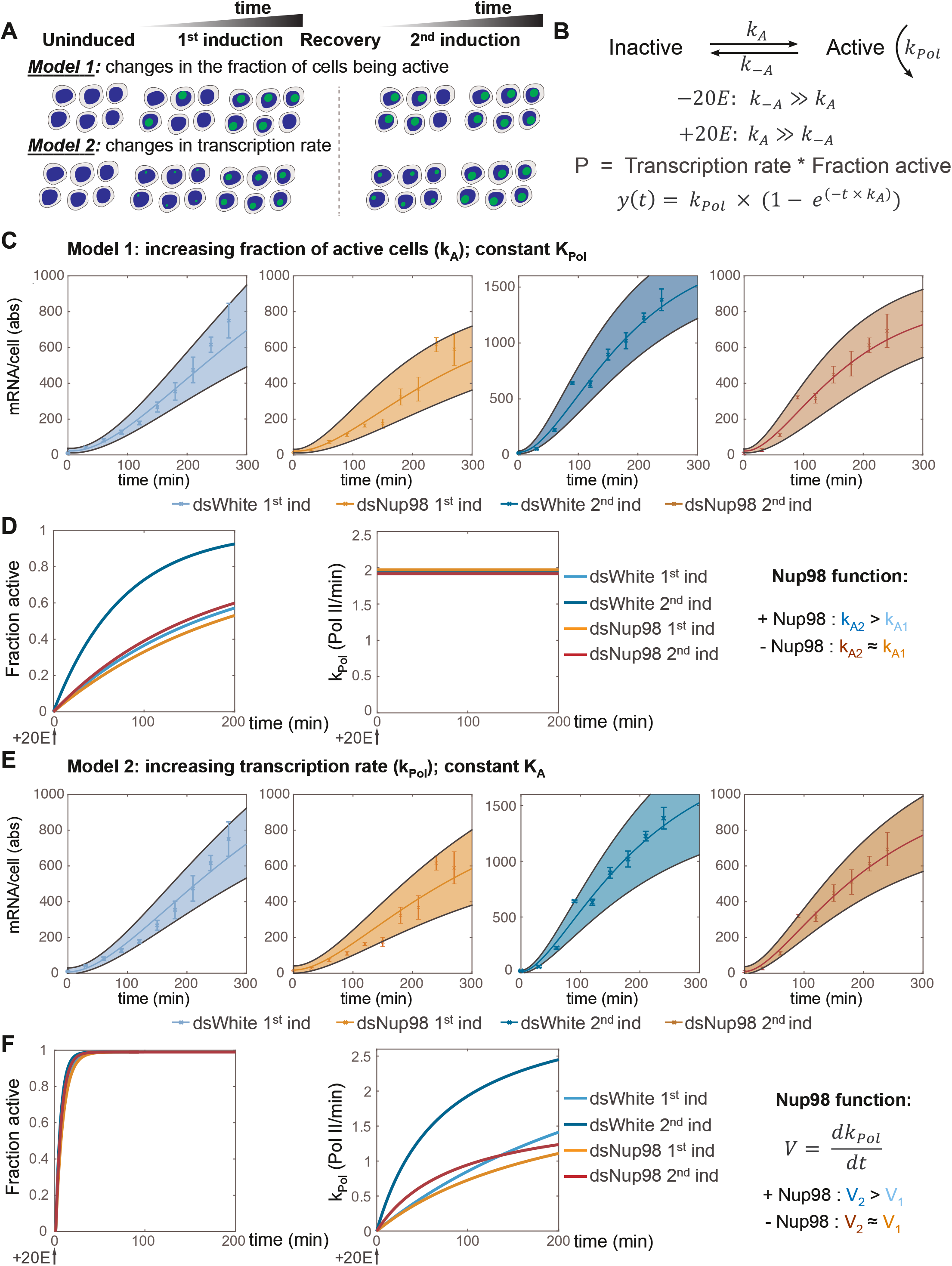
Modeling promoter state switching and increasing transcription rates using population-averaged qPCR data. **A.** A model utilizing two promoter states (active and inactive) contains two mechanisms that can account for the increase in the transcription rates observed by qPCR. Model 1: loci are slowly recruited into a transcriptionally active state upon first induction and more rapidly upon second induction. Once active, loci produce new mRNAs at a constant rate equivalent to the rate at which new RNA Pol II molecules enter productive elongation. Model 2: all cells are rapidly recruited into an active state, and the rate of transcription increases over time, slowly upon first induction and quickly upon second induction. **B**. Model describing the rate of recruitment into the active state k_A_, into the inactive state k_-A_, and the rate of recruiting RNA Pol II molecules into productive elongation k_Pol_. The presence of 20E permits k_A_ to be larger than k_-A_. The production rate as a function of time is given by the RNA Pol II recruitment rate multiplied by the fraction of active loci. In Model 1, k_A_ increases and k_Pol_ is constant during 20E exposure, whereas in Model 2, k_Pol_ increases and k_A_ is constant. **C**. Best fit and 95% confident intervals of Model 1 to qPCR data. **D**. Under Model 1, normal levels of *Nup98* are required to ensure k_A2_ > k_A1_, yielding a faster recruitment of loci into the active state (left) upon second induction, whereas k_Pol_ is constant under all conditions (2.0 +/- 0.2 RNA Pol min^-1^). **E**. Fit of Model 2 to data. **F**. Under Model 2, k_A_ is large and essentially identical between conditions (left), ensuring near-simultaneous activation of all loci (left). The role of Nup98 is to ensure a rapid increase in k_Pol_ upon second induction (right).

There are then two basic ways in which the mRNA production rate can increase over time in a two-state model. The most straightforward is by recruitment of loci into the transcriptionally active state, while the loading rate k_Pol_ is held constant (Figure 3A, Model 1 and Figure 3B). In this scenario, what would change over time is the fraction of cells in a transcriptionally active state (Figure 3A, Model 1). But once a locus enters an active state, mRNA production would occur at a constant rate. The increased transcriptional output during re-induction would thus stem from a higher fraction of cells being active in post-memory conditions. This scenario requires a k_A_ that is small such that any given locus is unlikely to enter the active state in any given minute. The gradual increase in the fraction of active loci would thus be proportional to the continuous increase in the average transcription rate (Figure 2E, right). Under Model 1, the role of Nup98 is to increase k_A_ such that a larger fraction of loci become active sooner.

We fit this model to the qPCR data. For each of the four scenarios (control/dsWhite or Nup98 knockdown, first or second induction), we found values and confidence intervals that described the accumulation curves (Figure 3C). Fitting each scenario independently, we found that k_Pol_ was roughly constant at 2.0 +/- 0.2 RNA Pol II per minute for all scenarios (average and standard deviation calculated across all four scenarios). Moreover, fit values of k_A_ are very similar for the three low expressing scenarios, control/dsWhite first induction and both inductions in Nup98 knockdown, at an average of 5.1 +/- 1.6 x 10^-3^ per minute (Figure Supplemental 3A). This means the curves representing the fraction of active loci as a function of time have a similar shape for these three scenarios (Figure 3D). In contrast, k_A_ for the second induction in control is 33 +/- 15 x10^-3^ per minute, a six-fold increase (Figure Supplemental 3A). In this model, Nup98 is required for this dramatic upregulation in rate of entry into the active state during the second induction.

Although Model 1 can explain increasing transcription rate at the population level, we can also envision an alternative scenario, where all cells enter an active state rapidly upon hormone stimulation, and what increases over time is the transcriptional rate at individual loci (Figure 3A, Model 2). The two-state model provides an additional means of regulation, in which k_Pol_ is not constant but instead increases with time (Figure 3A, Model 2 and Figure 3B). In the most extreme version of this scenario, k_A_ is very large such that all loci switch into the active state immediately upon hormone exposure. This would be consistent with the known rapid rate of nuclear import of hormone receptors, on the order of minutes (Nieva et al., 2007). In this scenario, the increasing transcription rate would result from increasing k_Pol_, and the increased rate of mRNA production during second induction would be explained by a quicker increase in transcriptional rate at individual loci. Under Model 2, the role of Nup98 would be to ensure a more rapid increase in k_Pol_ upon repeated hormone treatment.

To determine whether this scenario could explain our observations, we fit a model to our qPCR data in which the number of attempts that RNA Pol II makes to engage in transcription increases linearly with time. A natural limit to this rate is apparent when considering the physical size of the RNA Pol II holoenzyme. The minimum footprint of RNA Pol II on the DNA template is roughly 50nt (Krebs et al., 2017). The elongation rate in nucleotides transcribed per time divided by the RNA Pol II footprint determines the maximum RNA Pol II loading rate. The resulting effective RNA Pol II loading rate is therefore the inverse of the sum of the intervals for RNA Pol II loading and clearance. At fast attempt rates, the effective rate is limited by the clearance rate. In fitting this model, we used an elongation rate of 1500 nt/min (Izbans and Luseo, 1992) and allowed the footprint to vary as a free parameter. We additionally set k_A_ to be very fast (1000 per minute), thereby ensuring that nearly all loci become active within one minute of hormone exposure. With these constraints we were able to fit the qPCR data from each scenario to a model of increasing RNA Pol II attempt rate (Figures 3E, F) that described the accumulation trajectories. As before, three conditions (both control/dsWhite and Nup98-depleted first inductions and Nup98-depleted second induction) have similar acceleration constants, with an average of 11.7 +/- 2.2 x10^-3^ Pol II min^-2^. In contrast, control second induction has an acceleration constant that is about six-fold larger than the first at 65.7 x 10^-3^ Pol II min^-2^ (Supplemental Figure 3B). The physical size of the polymerase prevents the effective RNA Pol II loading rate from becoming indeterminately high, capping the value of k_Pol_ (Figure 3F) and permitting reasonable fit to data. In this model, the role of Nup98 is to increase the acceleration of the attempt rate k_Pol_.

Overall, both versions of the two-state model perform equally well at describing the qPCR data (Figure Supplemental 3C). Additionally, the two versions are not mutually exclusive; our qPCR data would be equally well fit by many possible combinations of values for k_A_ and k_Pol_ acceleration constants. To discern between these possibilities requires single-cell measurements of transcription provided by smFISH.

### 4. Single cell analysis does not support the two-state model

To discern between possible versions of the two-state model outlined above, we proceeded to determine whether they could account for the distribution of transcriptional activity observed in our smFISH analysis of *E74* (Figure 1). We determined the instantaneous transcriptional activity in individual cells by summing the fluorescence intensity of all nascent transcription sites in each cell and normalizing to the intensity of single mRNAs. The resulting values are the absolute amount of nascent RNA present at transcribing loci in units of the equivalent number of mature mRNAs (Little et al., 2013; Zoller et al., 2018). As mRNAs tend to be localized to the cytoplasm, the unit intensity is referred to as the “cytoplasmic unit” intensity (C.U.). The number of C.U.s associated with each transcribing site depends on the number of probe binding sites present, and thus on how many RNA Pol II molecules are present and how much of the transcript has been synthesized. The normalization is possible because our probes target only exon sequences and thus the fluorescence intensity is not affected by mRNA splicing or the presence of introns.

To determine if either of these two versions of the two-state model accounts for the actual transcriptional behavior in ecdysone-induced responses, we used the values for k_A_ and k_Pol_ from the fit to qPCR to simulate the distribution of C.U.s for both models at three time points (1, 2, and 4 hours post-induction) under all four induction conditions. Monte Carlo simulations were used to generate RNA Pol II positions for 100,000 cells under each condition and time point. The RNA Pol II positions were used in combination with the probe positions along the transcribed RNA to obtain a sum of probe binding sites present in the simulated nascent mRNA. This sum was then normalized to the number of probes present in a single mRNA to generate simulated distributions in units of C.U.s. Histograms of measured and simulated C.U. distributions were then compared and goodness-of-fit values calculated (Figure 4 and Supplemental Figure 4, and see Experimental Procedures).

**Figure 4.**
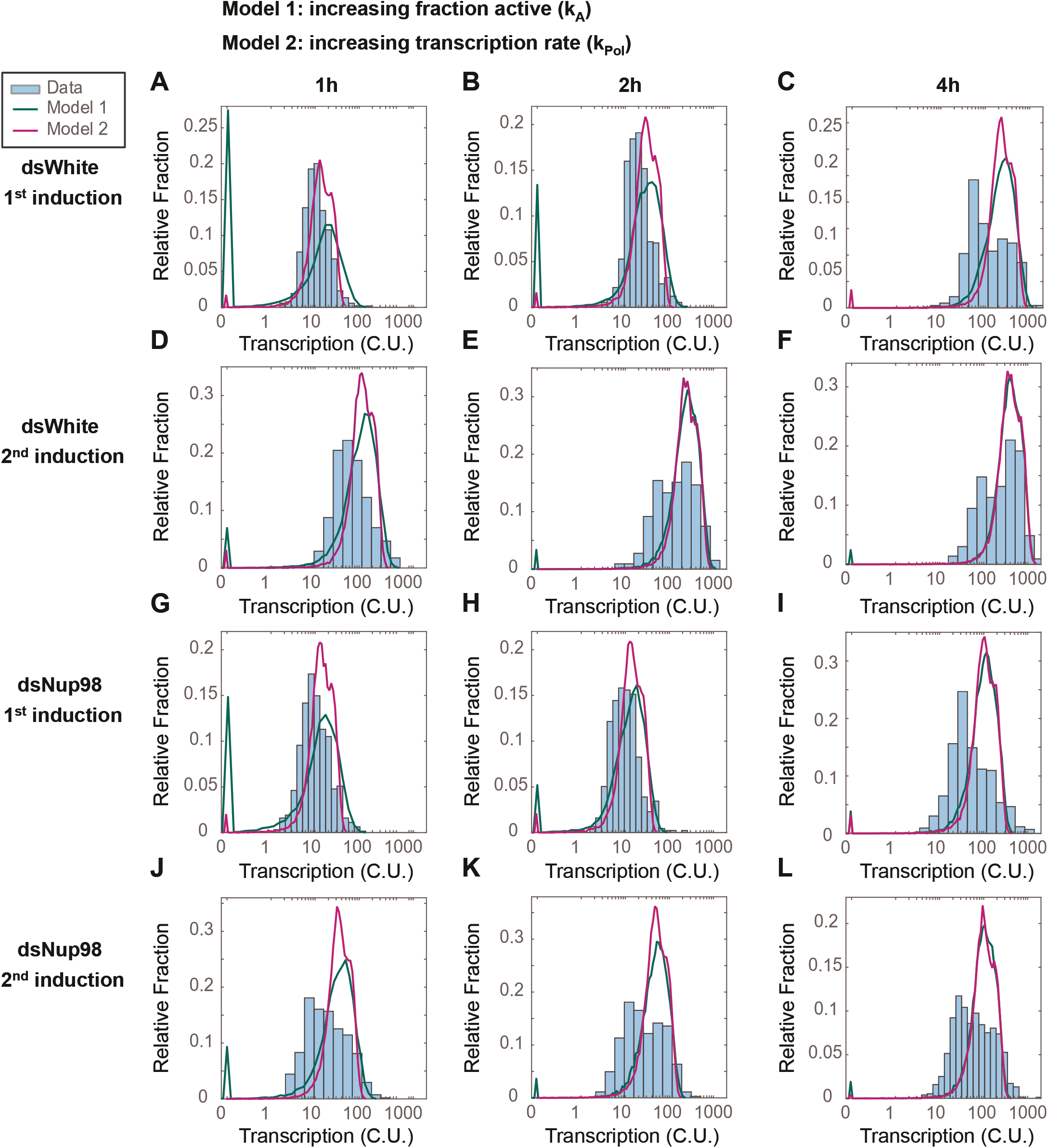
The two-state promoter model does not describe single-cell measurements of transcription. Histograms (cyan) show distribution of measured total instantaneous transcriptional activity in normalized units (C.U.), obtained from smFISH of *E74* as shown in Figure 1. Lines represent predicted values generated by simulation using best-fitting parameters for Model 1 (green) and Model 2 (magenta) under conditions of control (A-F) or Nup98 knockdown (G-L) conditions during the first (A,D,G,J), second (B,E,H,K), or fourth (C,F,I,L) hour of the first (A-C, G-I) or second (D-F, J-L) inductions.

We examined the distribution of transcriptional activity across hormone treatments and time points (cyan bars, Figure 4). Notable features in the observed distribution of transcriptional activity include a gradual shift in the activity levels as a function of time, as expected from the qPCR data, meaning that an increasing fraction of cells shift to transcribing at higher amounts. This shift occurs more rapidly for the control/dsWhite second induction (Figure 4D-F) than for other conditions. Moreover, during the inductions in control experiments, the histogram becomes broadly distributed even when fluorescence intensities are plotted on a logarithmic scale (Figure 4). The observed distribution begins to resemble a bimodal distribution in control cells at 4 hours (Figure 4F), demonstrating that a subset of nuclei have begun transcribing *E74* at rates much faster than earlier in the exposure, as measured in C.U.s. This is again consistent with the prediction from qPCR that the transcription rates in general are increasing. However, the distributions suggest that this increase occurs abruptly, not gradually, and that it occurs for only a subset of loci. Notably, this bimodal behavior is less evident in Nup98 knockdown conditions (compare Figure 4L to 4F). This suggests that Nup98 could be involved in a shift from low to high expression rates.

We compared the distribution of data to that predicted by the two-state models. Overall, the simulated distributions from either of the two two-state models provide a poor fit to data (Figure 4, compare simulated green and violet curves to cyan bars of observed data, and Supplemental Figure 4A). Model 1, with changing k_A_ and constant k_Pol_, consistently overestimates the fraction of cells with small numbers of transcripts at early times (green curves, Figures 4A, D, G, J, notice the early green “spike”). Model 2, with rapid k_A_ and increasing k_Pol_, suffers less from an overestimate of inactive cells. However, both models predict a narrowly distributed peak of highly active cells at four-hour time point in all four conditions. In contrast, measurements reveal broadly distributed expression levels, with many cells showing less activity than predicted (Figure 4C, F, I, L). Furthermore, hybrid models combining both more rapid k_A_ with more slowly increasing k_Pol_ suffer from both shortcomings in combination (data not shown). All intermediate values used in hybrid models suffer either from the overestimate of cells with zero activity at early times and/or the overestimate of transcriptional activity at later times. This stems from the failure of the two-state model to account for the large fraction of cells containing mRNAs in the first hour combined with an overestimate of narrowly distributed but highly active cells at late times. From the simulation results, we conclude that no two-state model correctly captures the transcription dynamics of the hormone response in either the first or second induction.

### 5. A four-state “memory switch” model explains single cell data of transcriptional memory

Given the distribution of transcriptional states we observed in our single-cell smFISH analysis (Figure 4), we reasoned that there may be two sub-populations of loci expressing *E74* at different rates, such that the loci are either in the low-or the high-expressing states. The slow emergence of the high-expressing state during exposure to ecdysone would be consistent with the gradual increase in transcription rate we have observed. Moreover, the rapid accumulation of transcripts upon hormone re-exposure suggests a mechanism for memory: loci that previously converted into the high-expressing state bypass the low-expressing state upon hormone re-exposure, and immediately enter the high-expressing state. We call such converted state the memory or high-expressing state, whereas loci that have not converted remain in a default low-expressing or non-memory state. Importantly, the model also suggests that the high expression observed during the second induction results from the continuous accumulation of loci into the memory state even after the withdrawal of hormone. In this case, the role of Nup98 may be both to ensure that converted loci retain information about their conversion state and to enact the continuous conversion of loci after hormone withdrawal. This would explain why the second induction strongly resembles the first induction when Nup98 is depleted: all loci reconvert to the default or non-memory state without Nup98 activity.

Based on this reasoning, we constructed a “memory switch” model containing four states: two non-expressing (non-memory and memory), and two expressing states (low- and high-expressing; Figure 5A, left). In this model, hormone exposure has two independent effects. First, similarly to the two-state model, hormone switches non-expressing loci into an expressing state at rate k_A_. As before, hormone ensures that the conversion to expression is irreversible as long as hormone is present. Second, hormone treatment converts loci from the default state (either non-memory or low-expressing) into the converted state (either memory or high-expressing). This conversion occurs at rate k_C_ and is necessarily slow. The two expressing states each have an associated rate of RNA Pol II loading, k_Pol_Low_ and k_Pol_High_. To account for the immediate entry of loci into the high expressing state upon second induction, the memory state is established irreversibly (and k_C_ is much greater than k_-C_ in normal conditions). Moreover, the model implies that once the hormone is applied, the conversion of loci to the memory state continues even after the hormone has been withdrawn, so that after the 24 hours recovery period in our hormone treatment regimen, most loci are found in the memory state. Under this model, the primary role of Nup98 would be to maintain a high k_C_, or the probability that cells remain in the memory or high-expressing state, such that depletion of Nup98 abrogates the maintenance of the memory state upon removal of hormone. Importantly, Nup98 depletion does not interfere with the conversion of loci into the high-expressing state. However, in the absence of Nup98, loci do not remember their prior exposure once hormone is withdrawn (Figure 5B, right), which explains why the second induction strongly resembles the first in Nup98-depleted cells.

**Figure 5.**
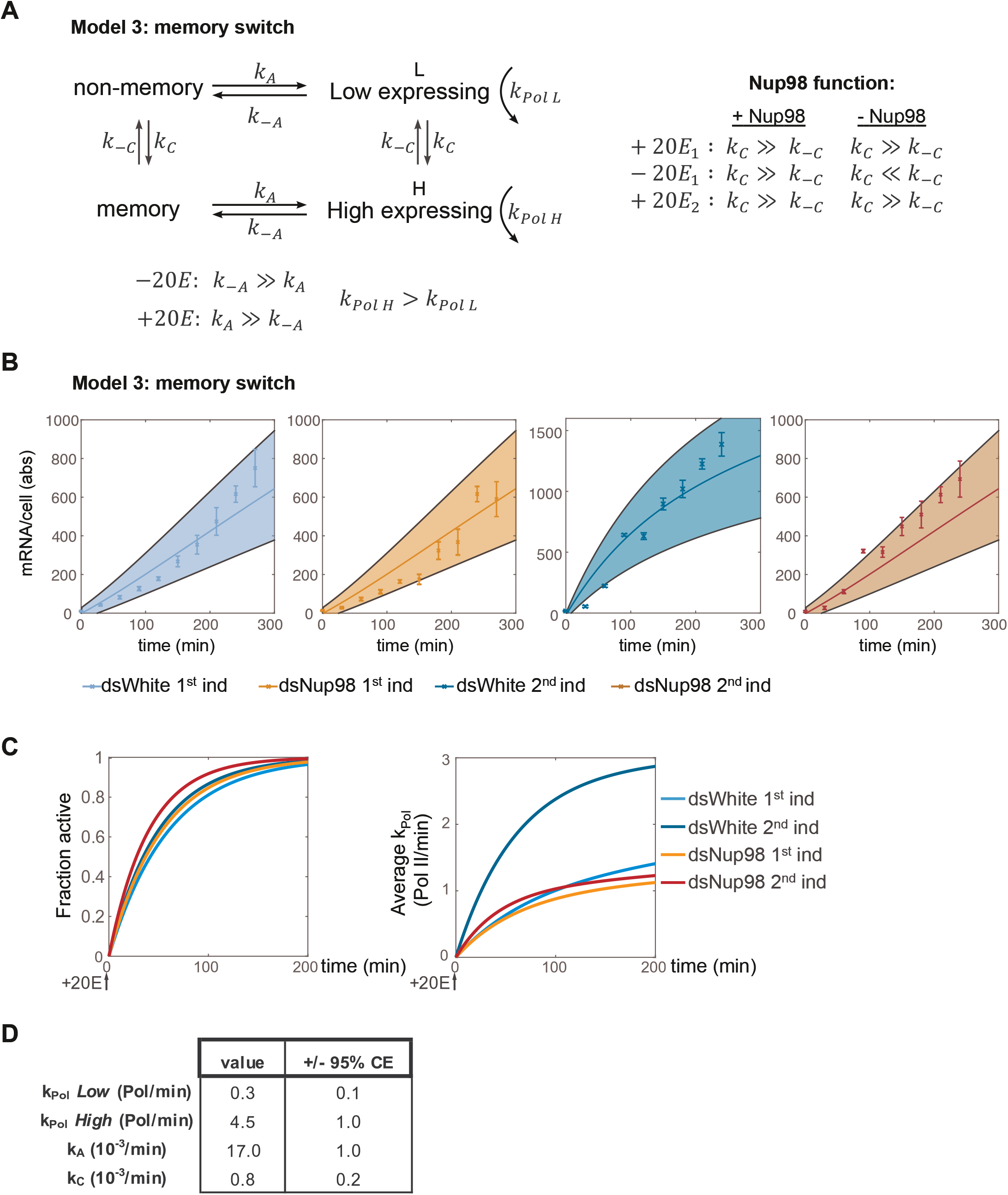
Fitting qPCR data to a four-state model of promoter memory. **A.** Promoters can occupy one of four states: low- and high-expressing states (L and H) with independent RNA Pol II rates k_PolL_ and k_PolH_; and two non-expressing states, non-memory (NM, the default state prior to 20E exposure) and memory (M). 20E has two roles: one, to activate transcription by ensuring k_A_ >> k_-A_, as in earlier models; and two, to increase the rate of conversion from NM (or L) into M (or H) by ensuring k_C_ >> k_-C_. The role of Nup98 is to maintain k_C_ >> k_-C_ upon withdrawal of 20E. **B**. Fitting of qPCR data to four-state model under each of four experimental conditions. **C**. The rate of recruitment of loci into the active state is similar between all conditions (left). Rapid mRNA accumulation upon second induction in control cells results from the rapid increase in average mRNA production rates. **D**. Best fit parameters with 95% confidence intervals.

If this model is correct, then a single set of values for the parameters k_C_, k_A_, k_Pol_Low_ and k_Pol_High_ should describe both the qPCR and smFISH results from all four conditions. To test this, we started by fitting the model to the qPCR data for each individual condition. We used the functions describing the average transcription rate over time (Figure 2E, right) as a constraint to narrow the range of allowable parameter values capable of recapitulating the functions. We then performed Gillespie simulations, searching a wide range of parameter values to find sets that reproduced the smFISH data. The resulting values captured the rise in the transcription rate and the resulting accumulation trajectories observed by qPCR under all four conditions (Figure 5B). As required by the model, the fraction of cells in either expressing state increased in a similar manner over time between all conditions (Figure 5C). This conversion into either expressing state occurs at essentially the same rate (on average, k_A_ = 17x +/- 1 10^-3^ per min, Figure 5D, see also Figure 5A). In contrast, the conversion into either the high expressing or memory state is very slow (k_C_ = 0.8 +/- 0.2 x10^-3^ per min) (Figure 5D, see also Figure 5A), meaning the characteristic decay time out of the default state is very long (>1200 minutes or about 20 hours). This means that the more time that passes after the first induction, the larger the fraction of loci that convert to the memory state, even after the withdrawal of hormone. In this manner, after the 24 hours recovery, the majority of loci (74%) are in the memory state (Supplemental Figure 5A), ready to enter the high-expressing state immediately upon re-exposure.

To verify the goodness of the model, we again performed Monte Carlo simulations and compared the resulting simulated distributions to smFISH data for *E74*. The simulation closely matches observation (Figure 6A) and provides a vastly improved fit relative to either of the two-state models (Figure 6B). The model thus successfully captures the emergence of the most highly-expressing cells during the first induction as a consequence of the slow conversion from low-to high-expressing state. Notably, the rate of RNA Pol II loading is 15 times greater for the high-expressing state (4.5 versus 0.3 RNA Pol II per minute) (Figure 5D) and approaches the maximum observed value found in *Drosophila* embryos (Little et al., 2013). The vastly increased expression rate during the second induction again requires Nup98 function; upon depletion, loci fail to re-enter the high expressing state upon re-exposure and are found in the default state upon re-induction (Supplemental Figure 5A).

**Figure 6.**
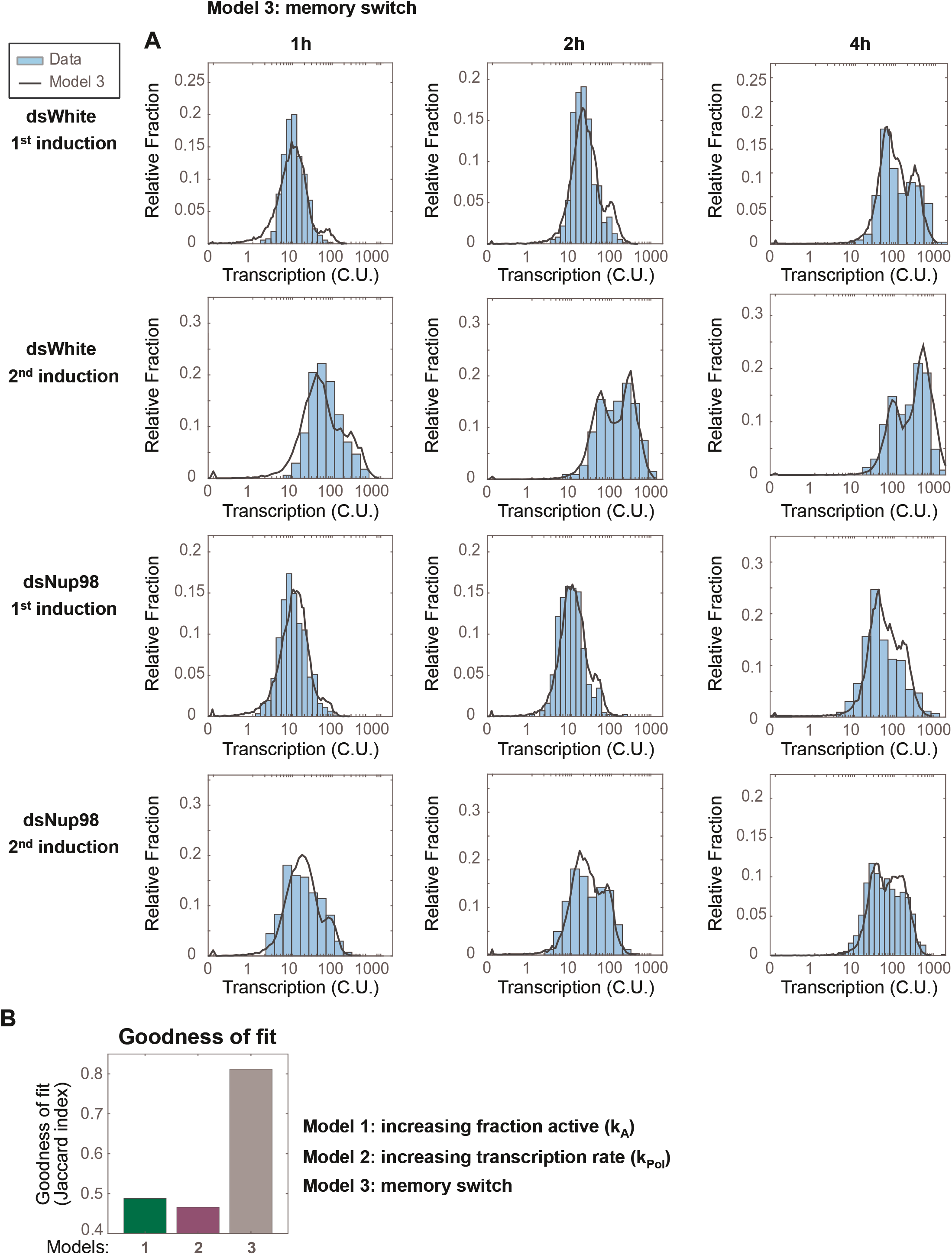
The memory model describes the distribution of transcriptional activity. **A.** Histograms (cyan) show distribution of measured total instantaneous transcriptional activity in normalized units (C.U.), obtained from smFISH of *E74* as shown in Figure 1. Lines represent predicted values generated by simulation using best-fitting parameters under the memory model. **B**. Goodness-of-fit is significantly improved for the memory model compared to either alternate models.

In summary, the four-state memory model encapsulates all the essential features of our observations. Our analysis puts forth a mathematical model that accurately describes the transcriptional memory behavior of ecdysone-inducible genes. The analysis predicts that in addition to activating default transcription, hormone exposure leads to a response event that drives a switch between non-memory and memory states.

### 6. The establishment of transcriptional memory is independent of transcriptional activity during the initial induction

The memory switch model, described above, reveals novel mechanistic insights into how transcriptional memory is established and maintained, and allows us to make predictions that can be tested experimentally. Thus, to further validate our model, we proceeded to test two of its main predictions. The most striking and surprising prediction of the model is the independence of k_C_ (the rate of conversion from low-expressing/non-memory to high-expressing/memory states) from k_A_ and k_pol_, which are rate constants that describe the transcriptional process itself. Otherwise stated, the model suggests that the ability of cells to establish the transcriptional memory response is independent from the process or the extent of transcription itself. The model predicts that at individual loci, addition of the steroid hormone ecdysone sets off the conversion into the high-expressing/memory state, but the parameters governing the rate of the conversion should be independent from the amount of transcription that takes place during the first induction.

We tested this prediction by two different approaches. First, we varied the amount of transcription by varying the length of exposure to 20E during the initial induction. We assessed the transcriptional memory response of the *E74* gene via RT-qPCR, using diminishing initial incubation times of 20E ranging from 10 min to 4 hours, after which cells were recovered for 24 hours and transcriptional memory was measured during re-induced conditions (Figure 7A). In agreement with the model, we observed that independently of hormone incubation times, cells responded with a similarly robust increase of transcription during the second induction. No significant changes were found in the *E74* transcriptional memory response between cells that were induced for 10 min versus 4 hours (Figure 7A), demonstrating that the memory state is established independently of the length of 20E incubation times and thus, of the length of time these loci were engaged in active transcription [18]. Second, to address this prediction, we utilized the transcriptional inhibitor FP to prevent transcriptional elongation altogether. Cells were treated with FP for 30 min before 20E induction. We monitored *E74* expression and observed no transcriptional activity in cells treated with FP during ecdysone induction, revealing the efficacy of FP blockage (Figure 7B). Cells were then washed out, and we measured the transcriptional memory response after 24 hours of recovery by RT-qPCR. Strikingly, we observed that blocking transcription during the first induction does not substantially affect the ability of the cells to have a robust memory response, indicating that the transition into the memory state is independent of mRNA production during the first induction (Figure 7B). There is a decrease of only 18% in the transcriptional memory response of FP treated cells at 4 hours of second induction, as normalized to the transcript levels of the house-keeping gene *rp49*, expression of which has been shown to be unaffected by ecdysone exposure (Shlyueva et al., 2014). Taken together, our experimental data support the prediction proposed by our modeling studies and highlight a previously unreported feature of transcriptional memory: that its establishment is independent from the length of initial stimulation and from the transcriptional activity derived from that activation process.

**Figure 7.**
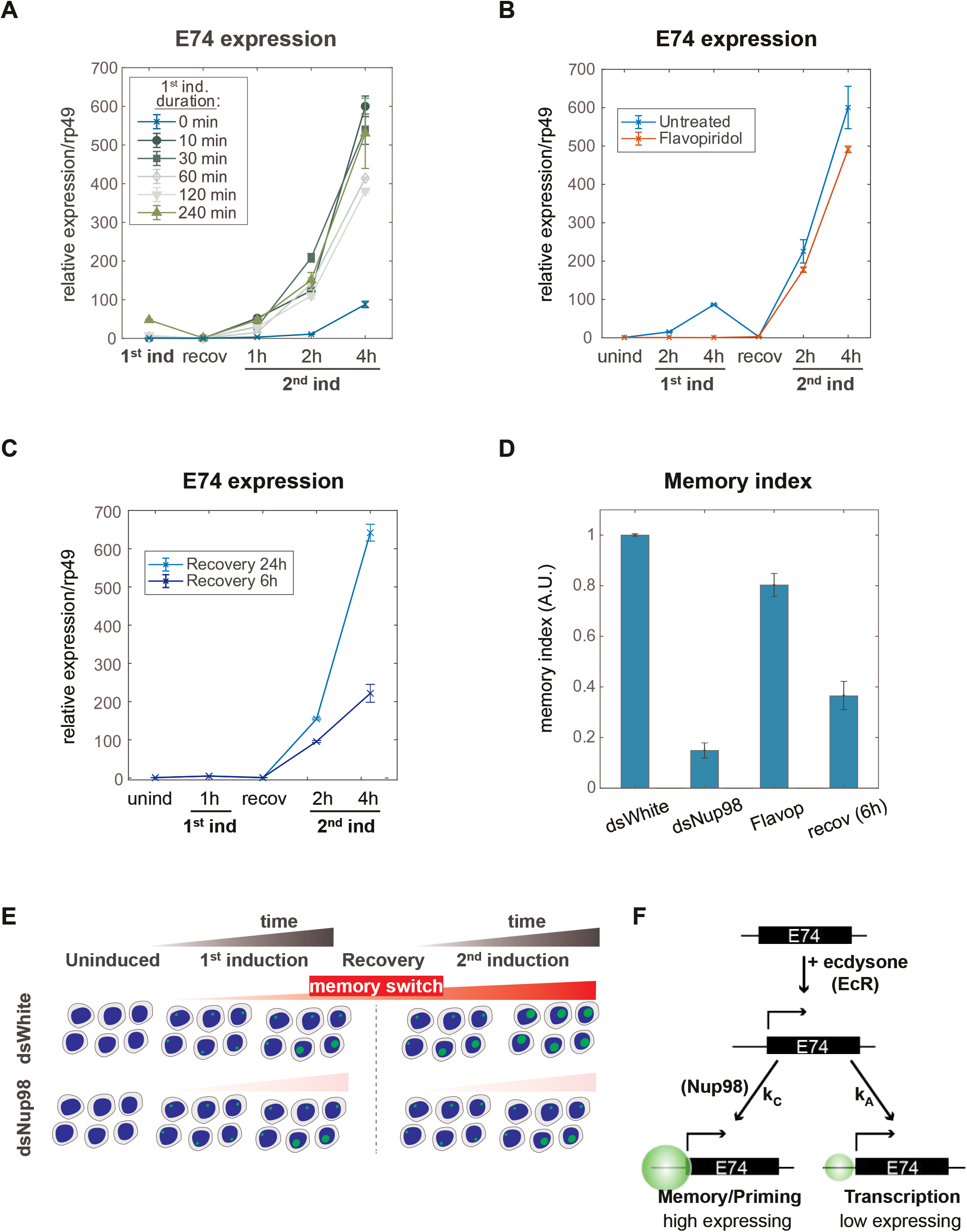
Induction of the memory state requires neither transcription nor prolonged 20E exposure. **A.** *E74* mRNAs measured by qPCR and normalized relative to *rp49* using different duration of 20E exposure during first induction. The data represent the mean of three independent experiments and the error bars the standard deviation of the mean. **B**. Flavopiridol inhibitor was added 30 min prior 20E first induction. After 4 hours, both the hormone and the transcriptional inhibitor were washed-out and cells recovered for 24 hours. 20E re-induced cells were collected and, *E74* expression was measured by qPCR from three independent experiments. Fold change values were normalized using *rp49* and error bars represent the standard deviation. **C**. *E74* mRNAs levels were measured by qPCR and normalized against *rp49* using two different recovery times (6 hours and 24 hours). The data represent the mean of three independent experiments +/- standard deviation. **D**. The Memory Index was calculated as the ratio of ecdysone induction’s slopes, and normalized relative to control (dsWhite). **E**. Memory switch model. Two events are triggered by exposure to 20E: loci rapidly enter into a low-expressing state and at the same time cells slowly switch from non-memory to memory. Memory is characterized by a high-expressing state, and importantly, cells continue to accumulate loci in the memory state after the hormone is withdrawn. When cells are exposed to 20E a second time, the converted memory loci transit into the high-expressing state and result into a more robust second expression. **F**. Implication of the memory switch model: upon hormone exposure, an activated locus engages in two separate activities, controlled by independent rate constants – entry into the default active state (controlled by k_A_) and transition into the memory state (controlled by k_C_). In the memory switch model, normal levels of Nup98 are required for cells to remain in the memory or high-expressing state (k_C_>>k_-C_), such that depletion of Nup98 abrogates the maintenance of the memory state upon removal of hormone.

The second prediction of the model that we aimed to test suggests that given the value of k_C_ derived from our modeling, the conversion from non-memory to memory state is relatively long, on the order of 20 hours (for a conversion of 62% of loci). We tested this timescale by shortening the recovery times after ecdysone induction. Cells were induced with 20E for 1 hour, and recovered for 6 or 24 before testing transcriptional memory responses. The model anticipates that cells that have been recovered for 6 hours should have a lowered memory response than cells that have been recovered for 24 hours. Consistently with the model, we found a reduced memory response for cells recovered for 6 hours (Figure 7C), underpinning the notion that transition into the memory state relies on mechanisms that have a relatively long timescale.

In order to compare the differences in transcriptional memory responses caused by distinct factors or treatments, we represented the observed differences in transcriptional memory responses as the “Memory index”, defined as the ratio of slopes in mRNA accumulation between first and second inductions, as measured by qPCR (Figure 7D). Comparing the Memory index among different analyzed treatments demonstrates that while depletion of Nup98 has a substantial impact on transcriptional memory, reducing Memory index by 85%, inhibition of transcription by FP affects transcriptional memory to a much lesser extent, reducing Memory index by only 18% on average, and shorter recovery time reduces Memory index by around 60% (Figure 7D). Together, these results support our computationally derived model of transcriptional memory and reinforce the notion that ecdysone-driven induction initiates two independent events at a given locus: 1) the rapid switch and transition into active transcription, and 2) the slow-acting Nup98-dependent conversion into the transcriptional memory state. The population dynamics of the transcriptional responses to ecdysone would thus reflect the changing mixture of the low-expressing and high-expressing/memory cells (Figure 7E), with Nup98 functioning as a key determinant for the prevailing fraction of high-expressing cells post-memory establishment. Our approach suggests that the main role of Nup98 in transcriptional regulation lies not with influencing transcriptional entry, but instead, with stabilizing and maintaining the specialized state of high transcriptional output, conversion to which is determined by a separate rate constant k_C_ (Figure 7F).

## Discussion

The different models by which the primed state of transcription is established and maintained have been primarily drawn from bulk cell population studies. Using the *E74* gene as a model, we aimed to understand the gain of transcriptional priming in single cells within a population. Together, our data show that the acquired *E74* transcriptional memory is characterized by a high transcriptional output from a sub-population of cells transitioning into a memory state. Our modeling and experimental approaches suggest the following model (Figure 7E-F). Upon ecdysone stimulation, two independent but simultaneous processes are initiated: 1) cells rapidly activate the ecdysone-dependent transcriptional response, characterized during this initial induction by a low rate of transcriptional output, and 2) concurrently, cells slowly progress into a specialized memory state defined by a high transcriptional rate, which is detected in subsequent ecdysone inductions. Importantly, we found that the transition into the memory state is independent of the transcriptional process stemming from ecdysone activation since blocking transcription with an RNA Pol II inhibitor does not compromise the memory response significantly. In agreement with this observation, different time intervals of transcriptional activation by hormone exposure do not deteriorate the cells’ ability to prime transcription, supporting the notion that the memory state is independent of initial levels of transcription. Another interesting conclusion from our model is that, if such a separation occurs generally at inducible genes, then phenotypes of certain epigenetic regulators may not show up until later on and would not be detected within a short time window, which is often used in Auxin-induced degradation system of rapid protein removal (Morawska and Ulrich, 2013; Nabet et al., 2018; Nishimura et al., 2009). As our findings reveal, on-going transcription and transcriptional memory may be controlled by separate mechanisms and would manifest their effects at different time scales.

A complex interplay of transcriptional factors, changes in histone modification and the incorporation of histone variants have been linked to the transcriptional memory response of multiple genes. Additionally, transcriptional memory in certain systems involves the cytoplasmic inheritance of a regulatory protein produced during the initial process of transcription, such that the transcriptional memory response is dependent on the protein’s level, which gets diluted in each cell division. This type of memory process has been found in yeast genes that respond to oxidative stress or changes in nutrients (Guan et al., 2012; Kundu and Peterson, 2010; Zacharioudakis et al., 2007). For example, in the case of the yeast *GAL1* gene, a key factor controlling the memory response is the passive inheritance of the trans-acting Gal1 protein itself, produced during transcriptional activation (Kundu and Peterson, 2010; Zacharioudakis et al., 2007). For the ecdysone-induced *E74* gene, this type of regulation seems unlikely since blocking all transcription still results in a robust memory response, demonstrating that the model of cytoplasmic inheritance described above does not extend to all memory systems.

Nup98 has emerged as an evolutionarily conserved factor required for transcriptional memory (Light et al., 2013; Pascual-Garcia et al., 2017). Our smFISH analysis, combined with mathematical modeling, revealed that active loci in a memory state initiate new transcription about 15 times faster than active non-memory loci. It also shed light on Nup98’s role in this process, demonstrating that Nup98 promotes the rate of conversion between non-memory to memory state, once the activating agent has been removed. Thus, Nup98 does not affect transcription during initial induction, yet cells are unable to remain in or transition into the memory state upon hormone withdrawal, causing a defect in the amplified transcriptional response upon secondary ecdysone stimulation. How does Nup98 promote the memory state in molecular terms? In line with Nup98’s role in securing the memory state, we have previously reported that Nup98 depletion disrupts the enhancer-promoter loops induced by ecdysone’s initial induction (Pascual-Garcia et al., 2017). Our current results support a model where the stabilization of enhancer-promoter contacts by Nup98 might form part of the memory state, supporting our previously proposed notion that changes in enhancer-promoter contacts are functionally separate from initial transcriptional activity. Consistently, we have also reported that Nup98 gains physical interactions with EcR and architectural proteins upon ecdysone stimulation, suggesting that they might also form part of the memory state (Pascual-Garcia et al., 2017). In agreement with the present data, the memory complex formed by interactions of Nup98, architectural proteins, and enhancer-promoter contacts were similarly found to persist during transcriptional shut-off. Overall our data are consistent with the idea that such architectural role of Nup98, as well as architectural functions suggested for other Nups (Ibarra and Hetzer, 2015; Kuhn and Capelson, 2018; Pascual-Garcia and Capelson, 2019; Sun et al., 2019), may play a role in memory maintenance. It is worth noting that at the galactose-induce yeast gene *HXK1*, where memory is associated with gene loop interactions between the promoter and 3’-end of the gene, the maintenance of these loops is mediated by an NPC component Mlp1 and is thought to promote faster recruitment of RNA Pol II due to retention of transcription factors in the loop scaffold (Tan-Wong et al., 2009). A similar mechanism may be executed by Nup98 in transcriptional memory of ecdysone-inducible genes, where a large complex, consisting of the looped gene, architectural proteins and key transcription factors, is “locked in” by Nup98 to create a high-expressing memory state. In this manner, it is tempting to speculate that phase-separating properties of Nup98 and other Nups may be involved in creating a complex specialized for high transcriptional outputs (Pascual-Garcia and Capelson, 2019; Schmidt and Görlich, 2016, 2015).

Another critical pathway implicated in transcriptional memory and in epigenetic memory in general is the deposition of histone modifications. H3K4 methylation is functionally linked to the transcriptional priming of a variety of genes (Ding et al., 2012; D’Urso et al., 2016; Jaskiewicz et al., 2011; Light et al., 2013, 2010; Muramoto et al., 2010; Sood et al., 2017), and more recently, histone H3K36me3 was correlated with the acquisition of memory upon IFNβ stimulation in mouse fibroblasts (Kamada et al., 2018). Although the interplay between transcriptional memory and deposition of histone modifications has not been fully explored in the *Drosophila* system, multiple connections between H3K4 histone methyltransferases (HMTs) and factors that regulate ecdysone-induced transcriptional kinetics have been reported. For example, HMT Trithorax-related (Trr), responsible for the deposition of H3K4Me1, has been found to regulate the transcriptional activation of ecdysone-inducible genes through interactions with EcR (Herz et al., 2012; Sedkov et al., 2003). Additionally, Nup98 has been found to interact with the related HMT Trithorax (Trx) physically and genetically (Pascual-Garcia et al., 2014; Xu et al., 2016), and to regulate some of Trx target genes during fly development (Pascual-Garcia et al., 2014). Trx is a well-known regulator of epigenetic memory during development, and is critical for maintaining the active transcriptional state of homeotic genes, which define tissue identity and correct body segments (Kingston and Tamkun, 2014). Whether the discovered role of Nup98 in maintaining a specialized active state plays a role in the epigenetic memory of Trx targets remains to be determined, but our findings open an intriguing possibility where transcriptional memory dynamics described here apply to the broader Trx-mediated memory.

Our modeling study indicates that the transition between non-memory to memory state is relatively long, lasting around 20 hours to convert 62% of loci (or 48 hours to convert 90% of loci) in an asynchronous cell population. In agreement with the model, our expression data show that reducing the time interval between ecdysone stimulations deteriorates the memory response. One possible explanation for such long-time scales of the k_C_ rate constant is that the formation of a memory complex requires stochastic or biochemical events that involve these time scales. Another explanation may be a communication between cell-cycle completion and *E74* memory. The transmission of transcriptional states from mother to daughter cells has been studied in multiple organisms from human cells to developing fly embryos and is thought to be critical for the maintenance of differentiation programs (Ferraro et al., 2016; Zhao et al., 2011). In *Drosophila* embryos, the use of live imaging to visualize transcription uncovered a 4-fold higher probability for rapid re-activation after mitosis when the mother cell experienced transcription (Ferraro et al., 2016). In mammalian cell culture, BRD4, a member of the Trx group (TrxG) proteins, has been implicated in faster re-activation kinetics of post-mitotic cells through the recognition of increased levels of H4K5ac on the previously activated transgene (Zhao et al., 2011). In both cases, the authors proposed that transcription of the DNA template might render it more susceptible to rapid re-activation after mitosis, which may be achieved through heritable changes of DNA-bound transcriptional factors, nucleosomes or histone modifications (Ferraro et al., 2016; Zhao et al., 2011). Additionally, it has been suggested that the process of mitosis itself is needed to help reactivate transcription to levels higher than those observed in the previous cell cycle, possibly via enhanced recruitment of regulatory factors to post-mitotic decondensing chromatin (Zhao et al., 2011). In this manner, it is plausible that the memory complex coordinated by Nup98 interplays with events immediately following mitosis to accelerate the dynamics of RNA synthesis, which could also explain why the enhanced transcriptional response depends on the length of time between ecdysone inductions.

## Acknowledgments

We are thankful to members of the Capelson, Little, DiNardo and Bashaw laboratories for insightful comments and suggestions. We are also grateful to Jessica Talamas (Dana Farber Cancer Institute) and Terra Kuhn (European Molecular Biology Laboratory) for critical reading of the manuscript and the Cell and Developmental Biology Microscopy Core Facility for confocal microscope use. M. C. is supported by the National Institutes of Health grant R01GM124143.

## Conflict of Interest

No competing interests are declared.

## Author contributions

P.P., S.C.L. and M.C. designed, interpreted experiments and wrote the manuscript. S.C.L. performed modeling studies. P.P. carried out all other experiments.

## Materials and methods

### Cell culture and chemicals

*Drosophila* embryonic S2 cells were grown at 25°C in Schneider’s medium (Gibco; 21720), supplemented with 10% (v/v) heat inactivated fetal bovine serum (Gibco; 10438034) and 1% (v/v) of penicillin-streptomycin antibiotics (10.000 U/mL) (Gibco; 15140163). For ecdysone induction experiments, 20-hydroxyecdysone (Sigma-Aldrich; H5142) was dissolved in 100% ethanol and used at 5 μM. Flavopiridol hydrochloride (Tocris Bioscience; 30-941-0) was prepared in 100% DMSO and added at 1 μM.

### dsRNA synthesis and transfection conditions

Double-stranded RNAs (dsRNA) fragments against *Nup98* or *white* genes were synthesized with Megascript T7 kit (Ambion; AM1334) using a PCR-ed DNA template from fly genomic DNA (primers listed in Supplemental Table 1). The integrity of the RNA was assessed by running the denatured product in a 1.2% agarose gel. To knock-down cells we mixed 10 μg of dsRNA per million cells with 7.5 μl of Fugene HD (Promega; E2311) in serum-free media and we incubated the dsRNA cocktail with exponentially growing cells for at least 3 days.

### RNA extraction and quantitative PCR

Total RNA was isolated in 1ml Trizol (Ambion; 15596018) and purified using Purelink RNA mini kit columns (Ambion; 12183018A) following manufactured instructions. RNA concentration was determined by measuring absorbance at 260 nm with a NanoDrop 2000 (ThermoFisher; ND-2000). cDNAs were synthesized using one-step RT-PCR kit (Qiagen; 205311). To amplify specific cDNAs, primers were designed to span an exon-exon junction (primers listed in Supplemental Table S1).

For absolute quantification analysis, the template for *E74* synthetic RNA standard was generated from a cDNA library and amplifying with specific primers covering the first two exons of *eip74ef-RA*. ssRNA was in vitro transcribed using the Megascript T7 kit. The concentration in grams per volume of the synthesized reference RNA was determined by fluorometric quantification using the RNA HS assay kit (ThermoFisher; Q32852) and Qubit 2.0 (ThermoFisher; Q32866). RNA dilution series were converted to copy numbers per volume using the molecular weight of the RNA standard (MW=126060 g/mole). For the RT reactions of the RNA standard dilutions we added an additional *eip74ef*-RA primer in addition to the random primers included in the reaction kit. To minimize our experimental error, we also constructed a DNA standard using *E74* specific primers. We used this standard to calculate the amplification efficiency of the *E74* primers by fitting a linear curve to log_2_(Ct) as a function of *E74* DNA template copy number; the resulting line contains a slope of - 3.566 and R^2^ of 0.9985. We constructed the RNA standard curve by single-parameter fit of a line of −3.566 slope to the previously retrotranscribed RNA dilution series. We also factored the losses of mRNA extraction and purification by adding a known amount of *in vitro* transcribed *E74* RNA to Trizol followed by purification and comparing the obtained amount of RNA; this approach resulted in 56% of RNA losses.

In all qPCR experiments, we used PowerSYBR Green PCR Master Mix (Applied Biosystems; 4367659) and QuantStudio 7 Flex thermal cycler (ThermoFisher; 4485701). Each qRT-PCR was repeated at least three times, the values were normalized to the *rp49* transcript unless otherwise stated, and the error bars represent the standard deviation of the mean.

### Single molecule RNA fluorescence in situ hybridization (smRNA FISH)

S2 cells were cytospun in poly-L-lysine treated coverslips and fixed with 4% para-formaldehyde for 10min. Coverslips were rinsed 3X in PBS and submerge in cold 70% EtOH for at least 24h. Complementary probes to the reading frame of *eip74ef-RA* were designed using Stellaris Probe Designer (https://www.biosearchtech.com/stellaris-designer), ordered from Biosearch and conjugated to Atto-565 (Sigma-Aldrich; 72464). Cells were washed twice with wash buffer [2X SSC, 10% formamide, 0.01% Tween-20] and equilibrated with hybridization buffer [2X SSC, 10% formamide, 10% dextran sulfate,1 μg/ml BSA]. The hybridization to probes was performed overnight at 37C in a moisturized chamber at a concentration of about 1 nM. After hybridization, samples were washed 3X in pre-warmed wash buffer and incubated at 37C for 30 min. We performed a final wash with 2X SSC, stained with Hoechst and mounted in ProLong gold (ThermoFisher; P36930). Imaging was performed by laser-scanning confocal microscopy on a Leica SP8 with a 63x oil immersion objective using identical scanning parameters and laser power for all samples. Voxel dimensions are 76×76×250 nm. The number of cells analyzed for each treatment were as follows, for the 0,1,2,4 hours times points - 1st induction, control: 1129, 803, 913, 1310; 2nd induction, control: 634, 954, 1420, 1351; 1st induction, dsNup98: 270, 759, 1413, 1068; 2nd induction, dsNup98: 643, 761, 780, 897.

### Image Analysis

#### mRNA detection and normalization

By scanning confocal imaging, RNA puncta corresponding to single mRNAs, RNPs, and sites of nascent transcription all appear as diffraction-limited objects whose fluorescence intensities correlate with mRNA content (Little et al., 2015, 2013, 2011). Puncta were separated from imaging noise and centroids of true diffraction-limited objects were found with difference-of-Gaussian (DoG) thresholding using custom MATLAB scripts (Little et al., 2013) on deconvolved images followed by fluorescence intensity measurements using raw images as described (Little et al., 2015). Objects were classified as mRNA puncta or nascent transcription sites on the basis of DoG intensities as described (Zoller et al., 2018). Both the numbers and fluorescence intensities of non-nascent puncta increase during hormone induction. To obtain measurements of mRNA content of each individual puncta in absolute units by smFISH, we adapted a prior normalization technique (Little et al., 2015, 2013, 2011; Zoller et al., 2018). Briefly, we divide the fluorescence intensity of all puncta by the mean intensity of the non-nascent site puncta found during uninduced conditions, since, in the absence of hormone, the mean number of mRNAs detected by smFISH corresponds to the mean number measured by qPCR. Individual puncta are thus assigned a value corresponding to the equivalent number of finished, mature mRNAs they each contain. Because non-nascent puncta are mostly found in the cytoplasm, we term the units of this measurement “cytoplasmic units” or C.U.s, the unit intensity of single mRNAs (Little et al., 2013). These measurements are thus in absolute units. When reporting transcriptional activity, we sum the fluorescence of all nascent site puncta assigned to each cell (Little et al., 2013; Zoller et al., 2018). This is advantageous because of the phenomenon of chromosome pairing, prevalent in Drosophila, in which homologous chromosomes are found in close physical association (Joyce et al., 2016). Pairing prevents unambiguous assignment of fluorescence to individual nascent sites, but does not preclude us from accurately assessing transcriptional activity in single cells. All transcriptional activity reported in Figures 4 and 6 is in units of C.U. per individual cell. In contrast, all parameter values derived from fitting are for individual loci, as described below.

#### Nuclear-cytoplasm segmentation

Hoechst stain was used to determine pixels corresponding to nuclear volumes as described (Petrovic et al., 2019), and the same approach was applied to the low-level nonspecific cytoplasmic fluorescence in the RNA channel to delineate total cell volumes. All RNA puncta were assigned to the nearest nucleus using on the basis of nearest-neighbor comparisons of the positions in 3D space of the centroids of puncta and nuclei, as described (Petrovic et al., 2019).

### Modeling

#### mRNA degradation

mRNA stability was assessed by collecting cells for qPCR at 30 min intervals following 4 hour of hormone induction followed by hormone removal and treatment with 1 μM flavopiridol to disrupt transcription. qPCR was used to measure *E74* levels relative to *rp49* as a function of time following the start of transcription inhibition. qPCR was performed in triplicate on cells treated for 72 hours with dsRNA against *Nup98* or *white* genes. The data were fit to a model of exponential decay using nonlinear regression to obtain mRNA degradation rates and 95% confidence reported in Figure 2B.

#### mRNA export

Image segmentation of cells into nuclear and total cell volume described above was used to assign all non-nascent RNA puncta to either the nucleus or cytoplasm based on the centroid positions of puncta in three dimensions. Puncta densities in both volumes were calculated as the number of puncta per cubic micron, and the fraction of puncta found in cytoplasm calculated for individual cells. Under an assumption of unchanging mRNA degradation, the ratio of cytoplasmic to nuclear densities is constant regardless of expression level as long as RNA processing and transport are rapid compared to mRNA degradation. Since degradation rates are slow and indistinguishable between Nup98 knockdown and control, we conclude that mRNA transport rates are unaffected by Nup98 depletion.

#### mRNA accumulation and production

Fitting of qPCR data was performed using nonlinear regression and the measured mRNA degradation rate to obtain 95% confidence intervals for parameter values. Fits were performed for each experimental condition individually to produce the values displayed in the panels. S2 cells are tetraploid with a division time of 24 hours, spending similar amounts of time in G1 and G2 (Cherbas and Gong, 2014). We therefore performed all qPCR fits using the assumption that the average number of *E74* loci per cell is six. All fit parameter values are reported in terms of individual loci. Curves shown in Figure 1B are piecewise polynomials. Models in Figure 2 each contain a single free parameter representing either the rate of constant mRNA production or the rate of linear increase in the mRNA production rate as a function of time. Both models in Figure 3 contain two free parameters. For the model of constant RNA Pol II loading, these are the first-order rate of conversion to the active state k_A_ and the rate of RNA Pol II loading while active k_Pol_. For the model of accelerating k_Pol_, k_A_ is fixed so as to virtually guarantee that all loci convert to the active state in 1 minute (as noted in the text, Figure 3F is plotted with a much smaller k_A_ for illustrative purposes). We introduced an additional parameter, the RNA Pol II footprint on the DNA template, to provide a natural limit on the maximum attainable k_Pol_.

To compare the fitting of the qPCR results to the measurements of transcriptional activity obtained by smFISH, the rates from fitting were utilized in Monte Carlo simulations of transcription (Gillespie, 1976). We simulated the time-dependent evolution of RNA Pol II numbers and positions on the *E74* gene for 100,000 cells for each of the four conditions (Nup98 knockdown or dsWhite/control, with or without 20E). The time scale of the simulation was set by the previously measured RNA Pol II elongation rate in S2 cells of 1500 nt/min (Ardehali and Lis, 2009; Buckley et al., 2014; Yao et al., 2007). The simulation was updated every 1/1500 min, the interval needed to transcribe one nucleotide. For the model of increasing production rate, the production rate was also updated based on the acceleration parameter multiplied by the elapsed time. RNA Pol II molecules and newly completed mRNA molecules were assumed to be evicted as soon as transcription was finished. For simplicity, we assumed that half of simulated cells are in G1 and half in G2, therefore containing either 4 or 8 *E74* loci. We therefore simulated 600,000 single loci and combined them randomly into 50,000 sets of 4 and 50,000 sets of 8 to represent 100,000 individual cells. We converted RNA Pol II positions into C.U.s by noting that the RNA Pol II position in the gene body determined how many probe binding sites are present in the nascent RNA. The total C.U.s per cell was attained by summing the number of probe binding sites present across all 4 or 8 simulated loci. Because all probes bind exonic sequences, neither the simulations nor observations are affected by splicing. With this procedure, we deduced parameter values in terms of single loci, rather than single cells. Starting conditions assigned to each cell were chosen at random from a Poisson distribution using the mean number of detected puncta under uninduced conditions. Simulated transcriptional activity values were taken at appropriate times after the start of the simulation (60, 120, and 240 minutes) to compare the simulated distribution to that observed by smFISH. Goodness-of-fit scores were calculated by finding the intersection of the areas under the normalized histograms of simulated and observed cells, using histograms generated with bins of identical width. For convenience, we compared log(C.U.) values due to the long tail of observed transcriptional activity, identical to a previously utilized approach for describing the distribution of mRNAs in ribonuclear protein complexes (Little et al., 2015). The areas of intersection between simulated and observed histograms for each of the three time points corresponding to each of four conditions were summed and then divided by the area of the union, generating an effective Jaccard index (JI) that was used as a goodness-of-fit score. A score of 1 indicates perfect overlap between observation and simulation. We note that the absolute value of the goodness-of-fit score for any individual model is not by itself informative and is heavily reliant on the positions of bin edges; in contrast, the comparison of scores between identically prepared distributions informs on the extent of difference in overlap between model and observation.

For the four-parameter model of transcriptional memory, we searched parameter space across eight orders of magnitude: For the first-order activation and conversion rates k_A_ and k_C_, between 10^-8^ and 1 min^-1^, and for the k_Pol_ rates, between 10^-7^ to 10 min^-1^. Fitting the curves derived from qPCR was uninformative, as a vast volume of parameter space can describe population-averaged data. We therefore performed Monte Carlo simulations of transcription in order to approximate the observed distributions of transcriptional activity at all 6 time points in control conditions. In principle, this entails a search through 4D space across cells simulated for 32 hours (4 hour 1^st^ induction, 24 hour recovery, and 4 hour 2^nd^ induction). However, the combination of the four parameter values is constrained to match the trend of the average mRNA production rate determined by the prior fit of the qPCR results. This effectively reduced the search space to three dimensions, since we could choose a value for one parameter and simulate across combinations of the remaining three. To reduce computational burden, we simulated 10,000 cells for each parameter set over the four hours of the first induction. We then inferred the fraction of loci that would be in the memory state with a given k_C_ following a 24 hour recovery period, and randomly assigned memory status to that fraction of loci, in order to avoid simulating the recovery period. We then simulated the 4 hour period of the second induction. The histograms of simulated and observed C.U.s were then scored by JI. This rapidly narrowed the possible range of parameter values to within approximately one order of magnitude. Subsequent fine-grained search of parameter space employed 100,000 cells per parameter set to find the maximum JI. The 95% confidence interval was assigned by bootstrap, taking 1000 random subsets of single cells from the data, redeploying the search of the narrow parameter window, and finding the mean and 2 standard deviations of the parameter values producing the maximum JI.

We note that this procedure requires that the transcriptionally active state has a lifetime significantly shorter than 24 hours after hormone withdrawal. We did not attempt to measure the lifetime of the active state. However, we note there is minimal transcription following 24 hours recovery, supporting an assumption of short active lifetime without hormone. Likewise, we did not attempt to measure the lifetime of the converted state, which would only become apparent in Nup98 knockdown cells upon hormone withdrawal. However, the fit parameters from control data provide a reasonable explanation of both control and Nup98 knockdown cells. This supports the hypothesis that the memory state lifetime is significantly less than 24 hours in the absence of Nup98, whereas memory is maintained indefinitely in normal cells and their progeny.

## Supplemental Figure Legends

**Supplemental Figure S1.**
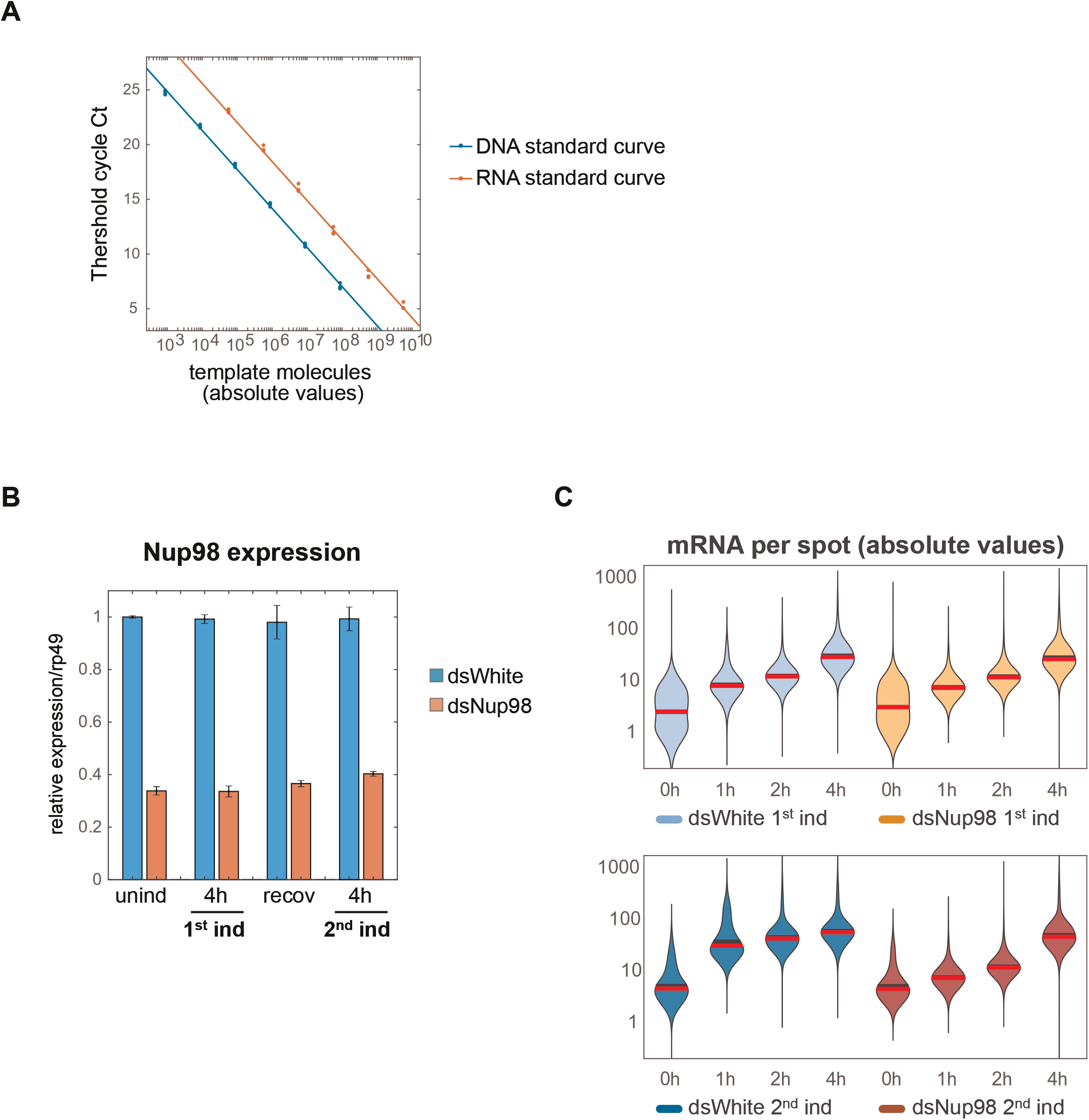
Absolute quantification of *E74* expression. **A.** Calibration of qPCR using known number of input DNA or in vitro transcribed RNA molecules. Ct (cycle threshold) refers to the number of cycles required for the fluorescent signal to cross the threshold at which all samples are compared. Note that the protocol is accurate across more than 6 orders of magnitude. **B**. *Nup98* expression was assessed by qPCR following the experimental conditions detailed in Figure 1B. qPCR data were normalized using *rp49* and error bars represent the standard deviation. **C**. Quantification of numbers of mRNA per spot detected by smFISH. Mean intensities of cytoplasmic puncta under uninduced conditions were used to normalize intensities of puncta under induction. Mean and median indicated by black and red horizontal lines.

**Supplemental Figure S2.**
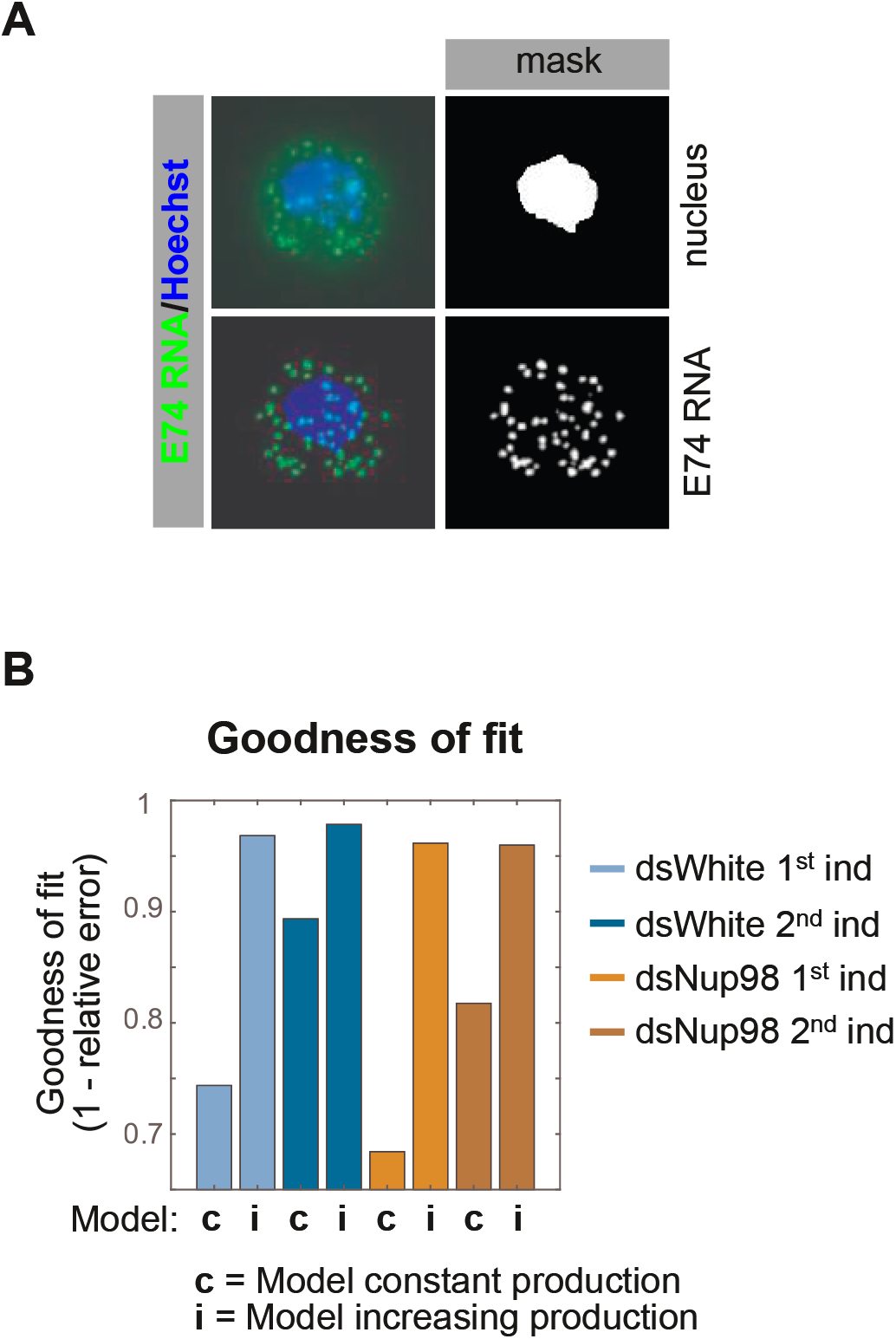
Models of mRNA export and production. **A.** Illustration of procedure to calculate mRNA export. Hoechst staining (blue in upper left) was used to generate a mask representing nuclear volume. Centroids for individual puncta were assessed to find the number of puncta assigned to nuclear and non-nuclear volumes. **B**. A model of constant production fails to describe *E74* accumulation. Goodness-of-fit was calculated as 1 minus the relative error, where the relative error is the mean of the absolute value of deviation of the simulated value minus the mean of the measurement at each time point divided by the mean of the measurement.

**Supplemental Figure S3.**
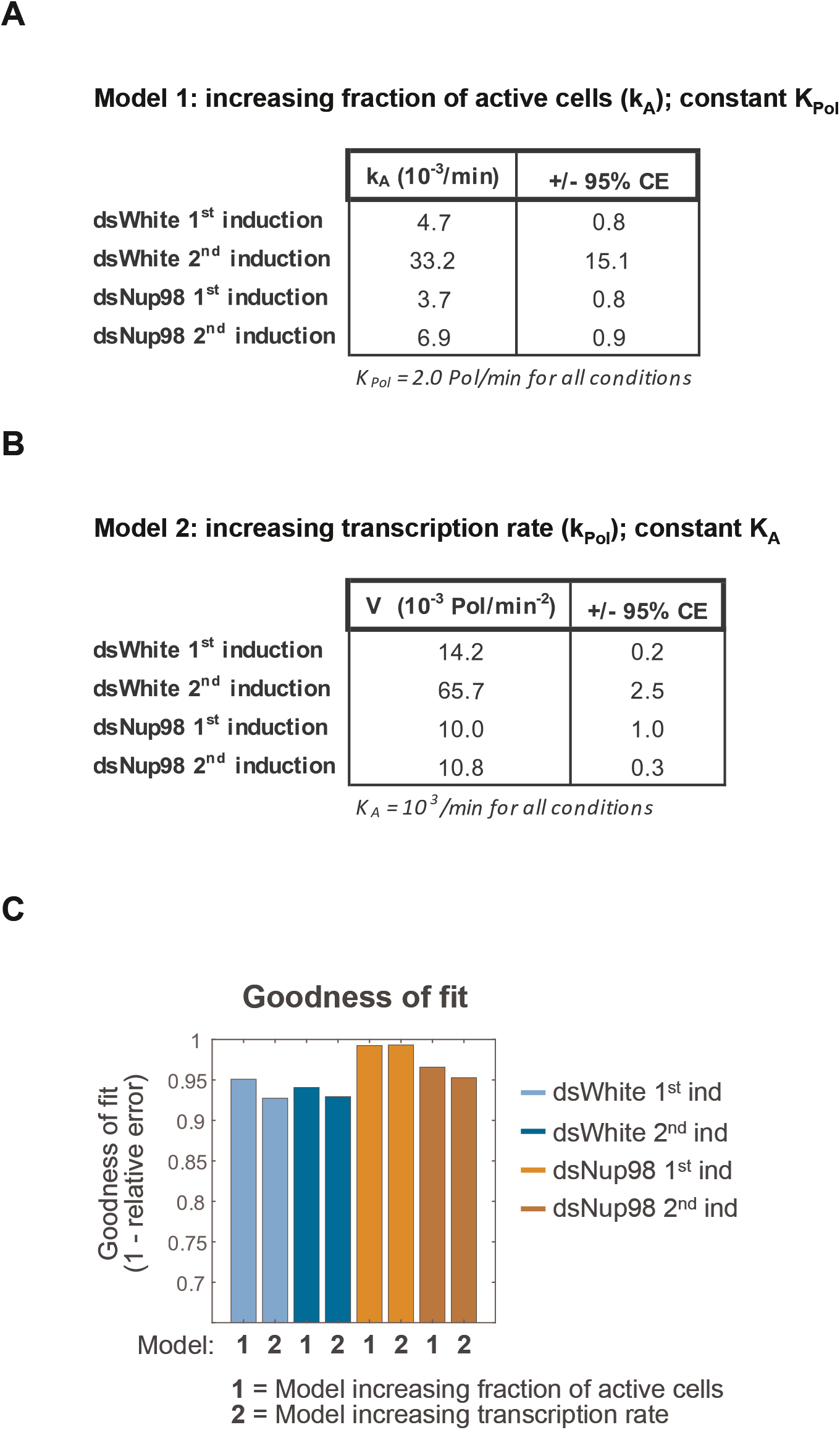
Fitting the two-state promoter model to qPCR data. **A.** Best-fitting values and 95% confidence intervals for a model of constant k_Pol_. **B**. Best-fitting acceleration rates describing the increase in k_Pol_ as a function of time. **C**. Goodness-of-fit values for each of the two mechanisms of increasing the transcription rate under an assumption that k_-A_ = 0. Values calculated as described in the legend to Fig. S2.

**Supplemental Figure S4.**
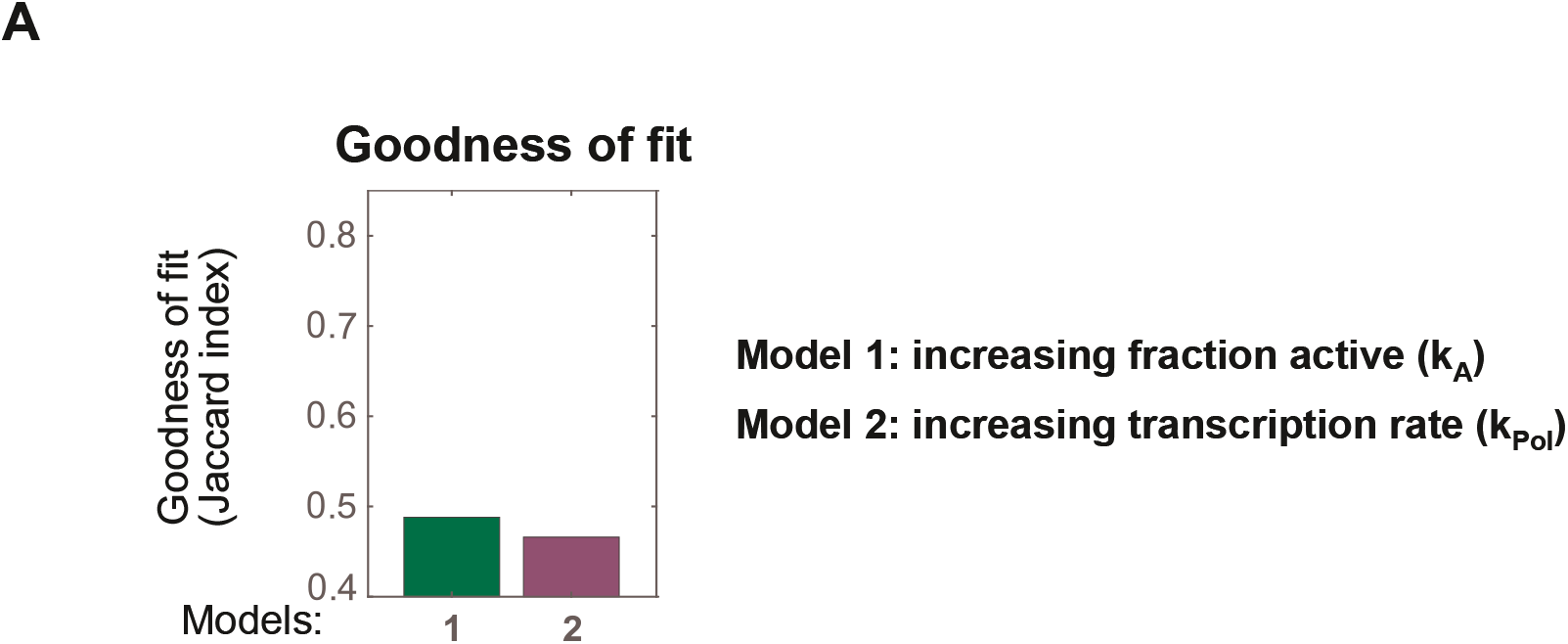
Goodness-of-fit for two-state model. **A.** Goodness-of-fit values for Models 1 and 2 describing the two-state model.

**Supplemental Figure S5.**
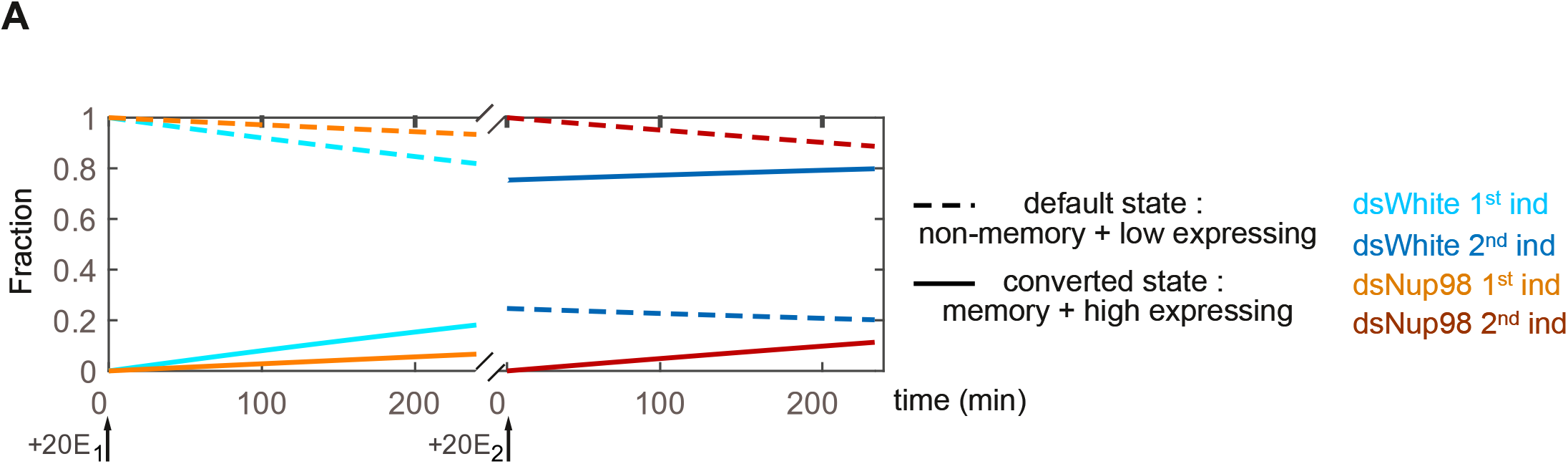
Memory model of *E74* expression. **A.** Fraction of loci in either the default state (dashed lines) or converted state (solid lines) as a function of time. In the presence of Nup98, loci continuously transition into the converted state starting immediately upon 20E exposure.

